# Multiple regulators control the biosynthesis of brasilicardin in *Nocardia terpenica*

**DOI:** 10.1101/2024.06.11.594307

**Authors:** Marcin Wolański, Michał Krawiec, Kay Nieselt, Tobias Schwarz, Dilek Dere, Bernhard Krismer, Carolina Cano-Prieto, Harald Gross, Jolanta Zakrzewska-Czerwińska

**Affiliations:** Faculty of Biotechnology, University of Wrocław, Wrocław, Poland; Institute for Bioinformatics and Medical Informatics, University of Tübingen, Tübingen, Germany; Interfaculty Institute for Microbiology and Infection Medicine Tübingen, Infection Biology Unit, University of Tübingen, Tübingen, Germany; Pharmaceutical Institute, Department of Pharmaceutical Biology, University of Tübingen, Tübingen, Germany

**Keywords:** immunosuppressant, actinomycetes, transcriptional factors, gene expression, secondary metabolite gene clusters

## Abstract

Brasilicardin A, BraA, is a secondary metabolite produced by the bacterium *Nocardia terpenica*, and a promising drug due to its potent immunosuppressive activity and low cytotoxicity. Currently, a semisynthetic approach confers production of a complete compound but suffers from insufficient heterologous biosynthesis of BraA intermediates used in the chemical semi-synthesis steps leading to only lab scale quantities of the compound. A better understanding of the involved gene expression regulatory pathways within the brasilicardin biosynthetic gene cluster, Bra-BGC, is a prerequisite to further improve production titers. However, the transcriptional regulation of the Bra-BGC has only been superficially analyzed, till now.

In this study, we comprehensively analyze the functions of several unstudied transcriptional regulators, KstR, SdpR and OmpR, encoded within the close vicinity of the Bra-BGC, and delve into the role of the previously described cluster-situated activator Bra12. We present, that Bra12 and the novel regulator SdpR, bind several DNA sequences located in the promoter regions of the genes essential for BraA biosynthesis. Subsequently, we demonstrate the complex regulatory network through which both regulators are capable of controlling activity of those gene promoters and thus gene expression in Bra-BGC. Furthermore, using the heterologous producer strain *Amycolatopsis japonicum*, we present, that Bra12 and SdpR regulators play opposite roles in brasilicardin congener biosynthesis. Finally, we propose a comprehensive model of multilevel gene expression regulation in Bra-BGC and propose the roles of locally encoded transcriptional regulators.

## 1. Introduction

Bacteria of the phylum Actinomycetota (formerly Actinobacteria) (Oren and Garrity, 2021) are prolific producers of clinically valuable secondary metabolites (SMs). These include a variety of compounds exhibiting diverse biological activities, e.g., antimicrobial, anticancer, and immunosuppressive. Among Actinomycetota, the genus *Streptomyces* is the main source of commercially available SMs, producing approximately two-thirds of antibiotics of natural origin. A tremendous increase in the number of sequenced bacterial genomes and the development of bioinformatics tools in recent years largely facilitated the identification of novel secondary metabolite gene clusters (SM-BGCs) – this has revealed that other “less-studied” than *Streptomyces* genera of Actinomycetota, e.g. *Amycolatopsis, Catenulispora*, *Nocardia*, can also serve as/represent precious sources of valuable metabolites (Ding *et al*., 2019; Engelbrecht *et al*., 2021). However, the discovery of novel compounds is hampered as most of the SM-BGCs are not expressed at all or only expressed at a very low level, making their putative products difficult to detect and to study under laboratory conditions. The genome mining strategy followed by genetic engineering tools and heterologous host expression allows for the unraveling of this treasure trove by enabling the activation of silent SM-BGCs (Gross, 2007; Rutledge and Challis, 2015; Ding *et al*., 2019; Lee *et al*., 2020; Bauman *et al*., 2021). One means of awakening silent SM-BGCs includes the engineering of transcriptional regulators (TRs) responsible for controlling gene expression in those clusters. This usually involves overexpression of positive and/or elimination of negative TRs. The choice of strategy often relies on a multistep experimental analysis of regulatory protein function. That includes identification of TR target promoters within the gene cluster, TR binding sites, and deciphering the TR mode of action, including putative interaction of the TR with small ligand compounds or/and other protein regulators, or/and its posttranslational modifications.

Brasilicardin A (BraA) is an immunosuppressive compound produced naturally by the human pathogenic bacterium *Nocardia terpenica* IFM 0406 (Shigemori *et al*., 1998). To date, the physiological or ecological role of BraA in its native host remained unknown. However, BraA is considered a promising lead in organ transplantation due to its high immunosuppressive activity and novel mode of action (Usui *et al*., 2006). Importantly, BraA exhibits lower toxicity than other drugs currently used, such as cyclosporin (Komaki *et al*., 2000). The biosynthetic production of BraA using the native producer strain is elaborate since it requires as a biosafety level 2 (BSL-2) classified strain strict safety measures during the production and workup phase and in addition, it exhibits low isolation yields. Furthermore, the preparation of this complex natural product by total synthesis (Anada *et al*., 2017; Yoshimura *et al*., 2018) has been impressively demonstrated but is economically not feasible and not sustainable. However, a recent semisynthetic strategy that employed chemical modifications of the BraE compound, an intermediate of BraA biosynthetic pathway obtained through heterologous expression in a *Streptomyces griseus* host, allowed for the economical production of BraA on a gram-scale (Botas *et al*., 2021). Despite these achievements, further improvement of the titer of BraE is desirable, as this may subsequently facilitate the generation of new brasilicardin derivatives (Niman *et al*., 2023). However, still little is known about the mechanisms that regulate the expression of the genes involved in brasilicardin biosynthesis, making this task challenging.

In *N. terpenica*, the brasilicardin biosynthetic gene cluster (Bra-BGC) involved in the synthesis of the core skeleton and BraA modifications comprises 13 genes (Fig. 2). These genes include *bra0*-*bra11*, which encode enzymes responsible for the biosynthesis of the compound, and *bra12*, which encodes Bra12, a crucial positive TR of the gene cluster. Bra12 is required for the transcription of all *bra0*-*bra12* genes (Schwarz *et al*., 2018a). However, the target promoter regions for Bra12 within the Bra-BGC have not been identified so far. Additionally, genes coding for other putative regulators, namely KstR, SdpR, LysRNt, and OmpR, were recently identified in the regions directly flanking Bra-BGC (see Tab. 1). Among them, only LysRNt has been studied to date (Wolański *et al*., 2021). LysRNt was shown to play a negative role in the biosynthesis of BraA intermediates, possibly by controlling the expression of a significant portion of the genes within the Bra-BGC through LysRNt binding within the promoter regions of the *bra0-1*, *bra12* genes and a putative promoter region of the *bra7* gene. Interestingly, it was also shown that the DNA-binding activity of LysRNt is subject to regulation via the intermediates of the BraA biosynthetic pathway.

The presence of multiple putative transcriptional regulators suggests a complex, and possibly influenced by external factors, regulation of Bra-BGC expression. Since these mechanisms remain elusive, we focused our efforts on collectively examining the roles of the unstudied regulatory protein candidates (KstR, SdpR, OmpR) and exploring the function of Bra12. In this study, we identify SdpR as a new player in the regulation of gene expression within the Bra-BGC. We demonstrate the possible interplay between SdpR and Bra12 regulators in controlling the activity of crucial *bra0-bra1* gene promoters.

## 2. Materials and Methods

### Bacterial strains, culture conditions

The strains used in this study are listed in Table S1 in Supplementary Information (SI). For *Escherichia coli* cultures the conditions, media, and antibiotic concentrations followed commonly used protocols (Sambrook and Russell, 2001); for *Amycolatopsis japonicum* cultures the growth conditions and media were the same as described previously (Schwarz *et al*., 2018a). Expression of recombinant proteins in *E. coli* hosts were performed as described in the paragraph protein purification.

### Cell cultures for RNA isolation

Seed cultures of *N. terpenica* IFM 0406 were grown from spore stocks in GPM medium (pH 7.0 ± 0.2) for 39.25 h at 37^◦^C and 150 rpm. Subsequently, four 300 mL Erlenmeyer flasks (A, B, C and D) containing 100 mL of GPM medium were inoculated with 300 μL of seed culture and incubated for 33 and 48 hours, respectively, at 37^◦^C with shaking at 150 rpm.

### RNA extraction

For RNA extraction bacteria were grown as described above. To prepare samples for RNA purification, 20 mL main culture aliquots from the four shake flasks A-D were quickly transferred after 33 h and 48 h, respectively, into 50 mL centrifuge tubes containing 2.2 mL of phenol-ethanol stop solution (one part phenol equilibrated with TRIS/EDTA at pH 8.0 and nine parts of EtOH). Upon mixing by vortexing, each solution was incubated for 5 min on ice and subsequently centrifuged for 15 min at 4.500 g and 4^◦^C. Supernatants were discarded, and the resulting *Nocardia* cell pellets were stored until RNA extraction at −80^◦^C. For cell lysis each pellet was suspended in 4 mL TRIzol and incubated for 5 min at RT. One mL of each TRIzol cell suspensions was transferred into 2 mL tubes with 0.5 mL zirconium beads (0.1 – 0.2 µm in diameter) and cells were disrupted by shaking employing a FastPrep-24 bead beating instrument (MP Biomedicals) 2 x 30 s at 6 m/s. The cell lysate was kept on ice for 3 min and subsequently mixed with 200 µL chloroform and centrifuged (5 min, 10.000 g at RT). For alcohol-based RNA precipitation, the upper aqueous phase was transferred into a new tube, mixed with 500 µL isopropanol and incubated 10 min at −80^◦^C. After centrifugation (10 min, 21.000 g, 4^◦^C), the supernatant was decanted and the resultant pellet was dried. The RNA pellet was dissolved in 100 µL RNase-free water and purified employing a NucleoSpin® RNA Clean-up kit according to the manufactureŕs protocol. The four extracted RNA samples (R1-33h, R2-33h, R1-48h and R2-48h) were finally obtained each in 50 µL RNase free water, quantified using a Nanodrop 1000 instrument, checked for quality using gel electrophoresis and stored at −80^◦^C.

### TEX treatment, cDNA library construction and sequencing

At first the four RNA samples were treated with T4 polynucleotide kinase. Subsequently, for the depletion of processed transcripts, equal amounts of each RNA sample were incubated with Terminator^TM^ 5’-phosphate-dependent Exonuclease (TEX) as well as mock treated without the TEX enzyme, as described previously (Sharma *et al*., 2010). To generate mainly sequencing reads of the 5’-end of the transcripts, the samples were not fragmented before cDNA synthesis. The cDNA libraries were constructed by Vertis Biotechnologie AG, Germany. For cDNA synthesis, the RNA samples were poly(A)-tailed to the 3’-end using poly(A) polymerase. Then, the 5’-triphosphate residues were removed using RNA 5’-pyrophosphatase. Accordingly, an RNA adapter was ligated to the 5’-monophosphate residue of the RNA. First strand cDNA synthesis was performed using an oligo(dT)-adapter primer and the M-MLV reverse transcriptase. The resultant cDNAs were PCR-amplified to approximately 10-20 ng/μL using DNA polymerase. Finally, the cDNAs were purified using an Agencourt AMPure XP kit and analyzed via capillary electrophoresis.

For Illumina NextSeq sequencing, the eight cDNA samples (R1-33h-TEX^+^, R2-33h-TEX^+^, R1-33h-TEX^-^, R2-33h-TEX^-^, R1-48h-TEX^+^, R2-48h-TEX^+^, R1-48h-TEX^-^, R2-48h-TEX^-^) were pooled in about equimolar amounts. Subsequently, the cDNA pool was eluted from a preparative agarose gel in the size range of 200-500 bp. The primers used for PCR amplification were designed for TruSeq sequencing according to the instructions of Illumina. The cDNA pool was single end sequenced on an Illumina NextSeq 500 system using a read length of 1×75 bp.

### DNA manipulations, plasmid and strain construction

For plasmid construction standard molecular biology procedures and the SLIC method were used (Kieser *et al*., 2000; Sambrook and Russell, 2001; Li and Elledge, 2007). Purification of plasmids and PCR products was performed using commercially available kits (Thermo Fisher Scientific and A&A Biotechnology). Transformation of *E. coli* cells with the linear or circular DNA constructs was conducted as described previously (Kieser *et al*., 2000; Sambrook and Russell, 2001). Plasmid constructs were verified by restriction digestion and/or DNA sequencing. To introduce plasmids into *Streptomyces* and *A. japonicum* hosts the original (Kieser *et al*., 2000) and the modified intergeneric conjugation procedure were used (Schwarz *et al*., 2018a). Plasmid and strain construction strategies are described in the SI. Plasmids, fosmids and oligonucleotides (supplied by Merck) used in the study are listed in Tables S1-S2; enzymes were purchased from Thermo Fisher Scientific and New England Biolabs.

### Protein purification

The recombinant KstR-His, SdpR-His, Bra12-His and OmpR-His proteins were purified using approaches described earlier in detail for LysRNtHis6 (Wolański *et al*., 2021). Briefly, the *E. coli* Rosetta^TM^ 2(DE3) strains harboring corresponding regulatory genes cloned in pET-21a(+) expression plasmid were used to express C-terminally His-tagged proteins. The proteins were purified on the Äkta Start system (GE Healthcare) using metal affinity chromatography resins (Talon column). The protein purity was assessed using sodium dodecyl sulfate-polyacrylamide gel electrophoresis (SDS-PAGE) (Laemmli, 1970) and staining (PageBlue^TM^ solution from Thermo Fisher Scientific).

### Protein-DNA interaction studies

Electrophoretic mobility shift assay (EMSA) and DNase I footprinting were used to investigate interactions between recombinant proteins and the DNA fragments representing either the selected regions on the BcaAB01 fosmid encompassing the gene cluster or the entire fosmid. For EMSA, a standard approach described earlier (Wolański *et al*., 2021) with a variety of modifications was applied – for details, see in the SI. DNase I footprinting was conducted as described in the previous study (Wolański *et al*., 2016).

### Detection of brasilicardin congeners

To assess the levels of brasilicardins production in *A. japonicum* strains, the procedure described in the previous study was applied (Schwarz *et al*., 2018a). Briefly, a preculture grown in tryptic soy broth medium (TSB) for 48 h was subsequently diluted with SM17 medium at a ration 1:100, and cultivated for another 72 h. The centrifuged supernatants were used directly for HPLC/MS analysis. The UV intensities obtained for each brasilicardin congener (BraC, BraC aglycon, BraD and BraD aglycon) were summed up to calculate total brasilicardin production.

### Bioinformatics tools, nucleotide and amino acid sequences

The web addresses for bioinformatics tools and other resources used in the study are listed in SI. Briefly, the following tools were used: protein parameters – ProtParam (Gasteiger *et al*., 2005); identification of protein domains - SMART (normal mode) (Schultz *et al*., 1998; Letunic and Bork, 2018) and CD search tool (default parameters) (Marchler-Bauer *et al*., 2017); searches for protein homologs - BLASTp (protein-protein blast); *in silico* prediction of protein DNA binding sites - MEME tool of MEME suite package ((Bailey *et al*., 2009) (search parameters were as follows: classic mode; maximum 3 different sequence motifs; occurrence 0 or 1 per sequence); generation of sequence logos – WebLogo (Crooks *et al*., 2004). The following sequence data are available on-line: *Nocardia terpenica* IFM 0406 genome sequence – EMBL/GenBank, accession number LWGR00000000.1 (Buchmann *et al*., 2016); the Bra-BGC sequence – GenBank, accession number MT247069. Direct links to the sequence resources are listed in the SI.

## 3. Results

Several previous attempts to achieve a high production yield of BraA in native and heterologous host systems turned out to be highly inefficient, most likely due to the low transcriptional activity of the Bra-BGC (Schwarz *et al*., 2018c; Schwarz *et al*., 2018a; Wolański *et al*., 2021). To verify this assumption, we examined global gene transcription in the native host *N. terpenica* IFM 0406 strain using RNA-seq. This analysis revealed that the expression of the brasilicardin gene cluster was indeed significantly lower compared to other randomly selected primary and secondary metabolism gene clusters (@Kay > e.g. gene cluster X, Y, Z) (Tab. S3AB).

Since our goal was to fully understand the mechanism regulating gene expression within the Bra-BGC, we embarked on a detailed study of the impacts of the transcriptional regulators (TRs) identified *in silico* within the Bra-BGC region (Wolański *et al*., 2021). In this study, we focused on Bra12 and three other putative regulators, namely KstR, SdpR, and OmpR, which have not been studied yet. To decipher their roles, we employed comprehensive experimental approaches following the general strategy presented in Fig. 1.

**Fig. 1.**
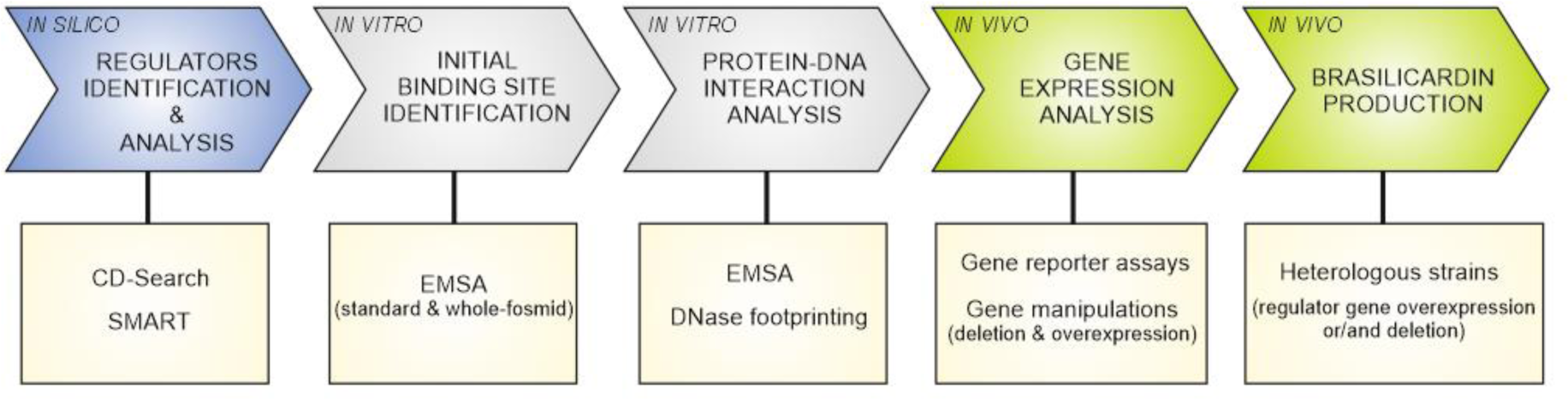
Overview of the experimental strategies used in this study. Schematic representation of the workflow used to decipher functions of transcriptional regulators identified within the Bra-BGC. The milestones are listed in the upper part of the panel, and the corresponding experimental techniques are shown below.

### 3.1. Local regulators bind targets within Bra-BGC

#### In silico analysis identifies putative novel regulators of Bra-BGC

To gain a first insight into the roles and confirm putative DNA-binding properties of the TRs identified in the Bra-BGC, we analyzed their amino acid sequences for the presence of functional protein domains using SMART and conserved domain-search (CD-search) tools. The analyzed proteins, including the previously studied LysRNt, comprise N-terminal DNA-binding domains (DBDs), except OmpR which contains the DBD at its C-terminus (Tab. S4 and Fig. S1A). The helix-turn-helix (HTH) (KstR and LysRNt) and winged-helix-turn-helix (wHTH) (SdpR, Bra12, OmpR) motifs are assumed to confer these DNA-binding functions. The C-terminal (N-terminal in case of OmpR) portions of those proteins encode putative regulatory domains (RD) involved presumably in signal-receiving functions, for instance, ligand binding (e.g., ADP in Bra12). The analyzed TRs are small proteins (MW < 35 kDa), except for Bra12 which exhibits a higher MW (66.1 kDa). The BLASTP search against nonredundant protein sequences in the NCBI database indicated that all TRs are relatively widespread in bacteria and, in the case of KstR, also in Archea (Tab. S4). A search against *N. terpenica* IFM0406 protein sequences indicated that each of the regulators has at least one putative paralogue protein in the native host (Tabs. S5A-E). Relatively large, in comparison to KstR and SdpR, number of paralogues in the case of the Bra12, OmpR, LysRNt (1-2 vs 14-27, respectively) suggests that the paralogs of the latter group of proteins are likely to cover various cellular processes and respond to a variety of environmental stimuli. Indeed, Bra12 homolog of AfsR/SARP family, AfsR, is a global regulator activated by phosphorylation and involved in complex regulatory networks in *Streptomyces* (Tanaka *et al*., 2007). Consistently, OmpR-family of regulators frequently comprise two-component systems involved in a transduction of environmental signals (Itou and Tanaka, 2001; Li *et al*., 2014). Importantly, *ompR* of Bra-BGC is accompanied by a divergently orientated gene (AWN90_RS33325) (Fig. 2) encoding a putative receptor histidine kinase that might be responsible for the phosphorylation of the OmpR regulator. The homologs of KstR and SdpR regulators are less extensively studied. The KstR belongs to a large family of TetR proteins that play roles in a variety of cellular processes (Ramos *et al*., 2005), including nitrogen metabolism (Jakoby *et al*., 2000; Loh *et al*., 2006). Interestingly, analysis of the gene organization indicates that *kstR* forms a presumable operon with two downstream genes *33470*-*33475*, coding for enzymes related with nitrogen (amino-acid) metabolism pathways. The SdpR putative regulator belongs to a large family of metalloregulators wide-spread in bacteria of which some act as metal sensing repressors involved in metal resistance processes, e.g. ArsR in *E. coli*, SmtB in *Synechococcus*, and CmtT in *Mycobacterium* (Erbe *et al*., 1995; Xu *et al*., 1996; Wang *et al*., 2005).

**Fig. 2.**
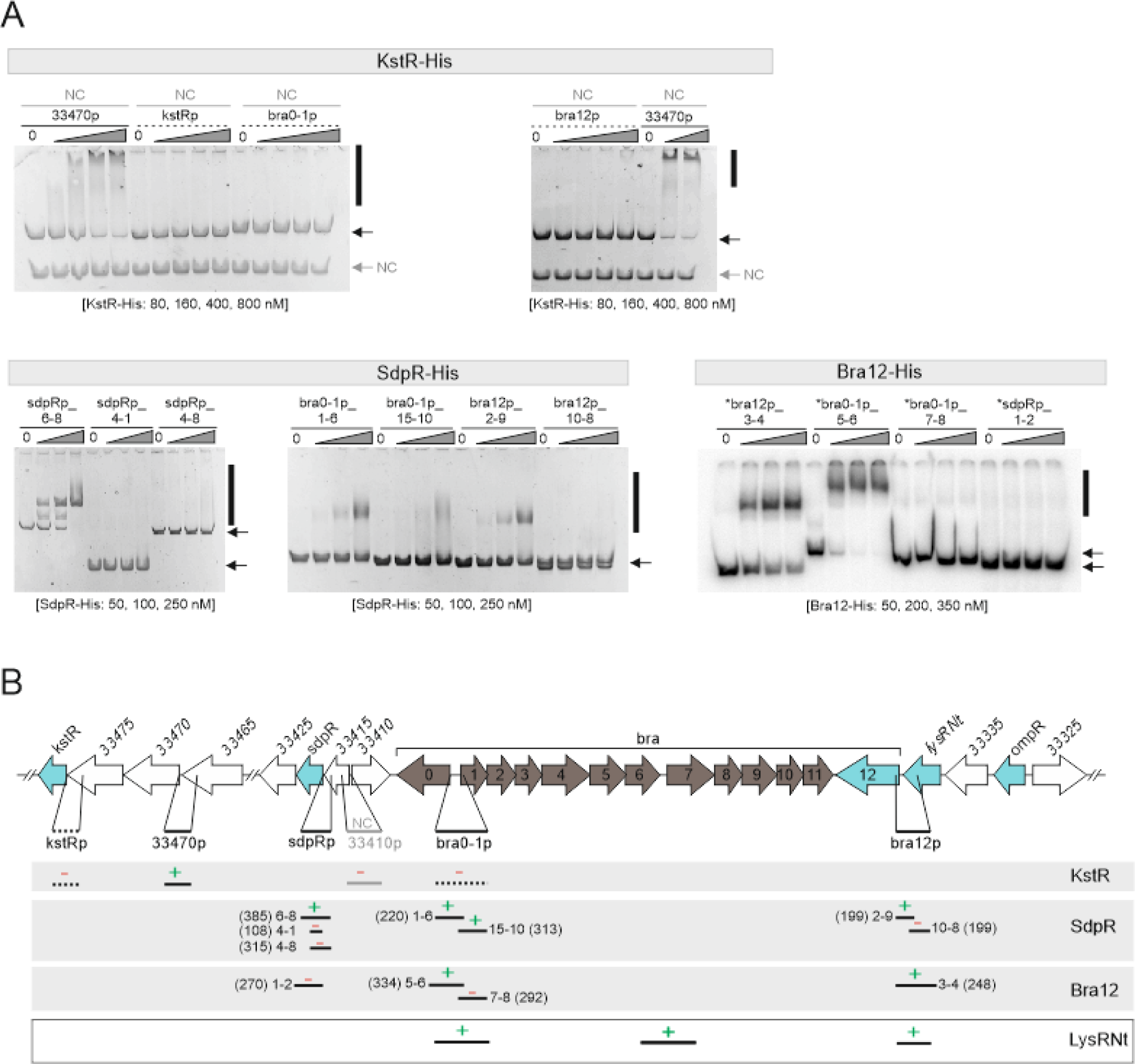
Identification of binding sites of regulatory proteins within the Bra-BGC. **(A)** EMSA. The recombinant proteins (Bra12-His, SdpR-His, KstR-His) at varying concentrations (indicated below the gels) were incubated with the same amounts of PCR-amplified DNA fragments comprising selected promoter regions (see Fig. S2-5). Unlabeled DNA was used for assays with KstR and SdpR proteins, and ^32^P-labeled DNA (marked with an asterisk) for assays with Bra12. The DNA fragments indicated with the solid lines represent regions identified in the preliminary EMSA (Fig. S2-5), those in gray indicate the internal negative control (NC) (33410p fragment). Black vertical bars along the gels indicate observed protein-DNA complexes; black and gray arrows indicate unbound DNA fragments and the internal NC. **(B)** EMSA - graphical summary. The DNA fragments analyzed are indicated by short horizontal bars. Numbers next to those bars correspond to primers used in the PCR reactions (Tab. S2); numbers in brackets represent fragment sizes (bp). The presence and absence of protein-DNA interaction is indicated by ‘**+**’ and ‘**–**’ symbols, respectively. The very bottom panel presents previously identified binding locations for the LysRNt protein (Wolański et al., 2021a).

#### Regulatory proteins bind to several promoter regions within Bra-BGC

To identify the genes putatively controlled by the regulators, we sought for the regulatory protein binding sites within the Bra-BGC using an *in vitro* approach based on electrophoretic mobility shift assays (EMSAs). Proteins used in these assays were expressed as C-terminal fusions with Hisx6-tag (Fig. S1B) and purified as previously described (Wolański *et al*., 2021). In the initial screen for the TR binding sites, we used as the DNA bait the restriction digested fosmid bcaAB01 that comprises the Bra-BGC and the cluster flanking regions. This approach led to the identification of several DNA regions bound by the KstR, SdpR, and Bra12 proteins over bcaAB01 (Figs. S2-4). However, the screen turned out to be unsuccessful for OmpR as we did not observe an interaction of this protein with any bcaAB01 fosmid DNA fragment (Figs. S5). This resulted in the exclusion of the protein from further analysis.

Next, to more precisely examine if the other regulators bind to regions containing gene promoters (designated using RNA-seq results, Fig. S11), we performed a set of EMSA experiments using promoter regions located within the fished-out DNA fragments. These included promoters of the investigated regulatory genes *sdpR*, *bra12*, and *kstR* (in the latter case, the *33470* gene promoter was used as the *33470*-*33475*-*kstR* form putative gene operon), and essential biosynthetic genes *bra0*, *bra1* (sharing a common divergent intergenic promoter region), and *bra7* (forming the presumed *bra7*-*bra11* sub-operon) (Wolański *et al*., 2021). The EMSAs revealed interaction of both SdpR and Bra12 regulators with the fragments comprising the divergent promoter region of *bra0*-*bra1* genes (bra0-1p) and the promoter region of *bra12* gene (bra12p) (Fig. 2 AB). In addition, SdpR exhibited binding to its gene promoter region (sdpRp). Noteworthy, the pattern observed for the SdpR-sdpRp interaction suggested the presence of multiple SdpR binding sites within the investigated promoter region. The KstR protein showed interaction with the 33470p fragment, which contained a presumed promoter region of the putative *33470*-*33475*-*kstR* gene operon. However, since *33470*-*33475* genes presumably do not play a role in BraA biosynthesis, as inferred from previous studies (Schwarz *et al*., 2018a), we excluded the KstR from further analysis at this stage. Importantly, the putative promoter of *bra7* was not bound by either SdpR or KstR (Fig. S6). This region was previously demonstrated as the target for another Bra-BGC regulator, LysRNt (Wolański *et al*., 2021).

In sum, the *in vitro*-based protein-DNA interaction studies identifed SdpR and Bra12, as the most promising regulators for further analyses. Both SdpR and Bra12 proteins bind to the promoter regions of *bra0-bra1* and *bra12* genes crucial for brasilicardin biosynthesis. Interaction of SdpR with the *sdpR* gene promoter region suggests that SdpR presumably regulates transcription of its own gene.

### 3.2. SdpR and Bra12 regulate the activities of gene promoters within the Bra-BGC

#### SdpR and Bra12 bind specific sequences within the promoter regions

EMSA analyses revealed *sdpR*, *bra0-1*, and *bra12* gene promoter regions as the targets for SdpR and Bra12 regulators in the Bra-BGC. To gain more insight into protein-DNA interactions, we next aimed to identify the binding sites and, furthermore, to determine the binding sequences for both the TRs.

First, we examined SdpR binding sites using the DNase I footprinting and EMSA. The footprinting assay carried out in the presence of the DNA fragment that included the promoter of the *sdpR* gene (sdpRp_1-2) revealed two DNase I-protected regions (Fig. 3AB), indicating the presence of two SdpR binding sites. This finding was in line with the previous assumption (Fig. 2A) and was also confirmed using shift assays with the subfragments of the *sdpRp* region (Fig. S7A). Interestingly, in the DNase I footprinting, we also observed the presence of the DNA region prone to DNase I digestion (marked with asterisks in Fig. 3AB). This DNase I hypersensitive site might result from protein induced DNA bending occurring when a protein interacts with a DNA region containing closely located binding sites. Importantly, we were unable to observe any DNase I-protected regions while investigating the interactions of SdpR with the fragments that encompass *bra0-1* and *bra12* promoters (data not shown). This can be explained by a much weaker interaction of SdpR with those two fragments (weak interaction at ∼100 nM of SdpR) than with the fragment comprising the *sdpRp* region (strong binding at 50 nM of SdpR) as observed earlier in EMSA experiments (Fig. 2A). Thus, to narrow down the DNA sequences that contain SdpR binding sites within these regions, we used standard and competition EMSA (Fig. S7B-C). These results exhibited binding of the SdpR regulator to two distinct locations within *bra0-1p*, and to one location within the *bra12p* region. The sequences of the identified fragments and the two *sdpRp* subregions comprising SdpR binding sites were further subjected to an *in silico* search for common sequence motifs using the MEME tool (Fig. 3C, left panel, and Fig. S7). This led to the discovery of a 14-nucleotide SdpR binding sequence, hereafter referred to as the SdpR box (Fig. 3C, right panel). The double SdpR boxes have been identified within the *sdpRp* and *bra0-1p* regions, and a single SdpR box within the *bra12* promoter region. The boxes exhibit a unique arrangement and locations with respect to the corresponding transcriptional start sites (TSSs) in the corresponding gene promoter regions (Fig. S8 and Fig. 7, Tab. S6).

**Fig. 3.**
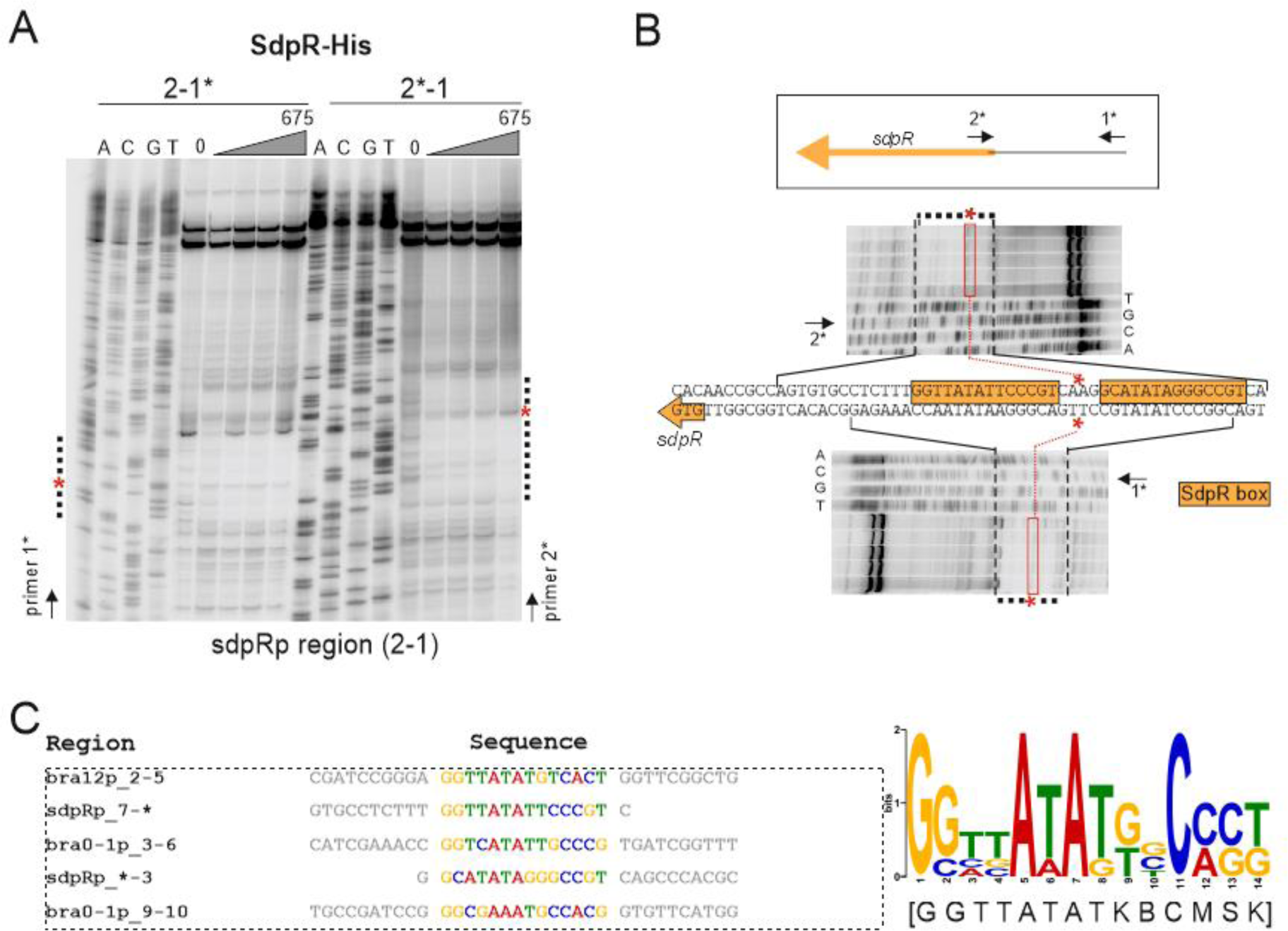
Determination of the SdpR binding sequence. **(A)** DNase I footprinting. The SdpR-His protein (25, 75, 225 and 675 nM final concentrations) was incubated with the ^32^P-labeled sdpRp_1-2 DNA fragment (270 bp) (marked with the asterisk), followed by DNase I treatment. The DNase I-protected or DNase I hypersensitive regions are marked with vertical dotted lines and red asterisks (*), respectively. The sequencing reactions are designated with A, C, G and T. **(B)** The detailed analysis of DNase I protected sequences and the locations of SdpR boxes. **(C)** (left). The alignment of the sequences containing the predicted SdpR boxes. The MEME *in silico* tool was used to predict SdpR sequences within the DNA fragments selected by EMSA (see Fig. S7). (right) The SdpR box sequence logo and consensus sequence (below in square brackets) were obtained using MEME.

Next, we aimed to identify Bra12 regulator binding sites within the *bra0-1p* and *bra12p* regions. As no DNase I protection patterns were observed in the footprinting assays (data not shown), we applied as previously an EMSA (including competition assay) approach. In the initial EMSA experiments, we narrowed down the DNA fragments comprising Bra12 binding sites in both promoter regions (Fig. 4AD). Moreover, the results also suggested the existence of single and double binding sites within the *bra12p* and *bra0-1p* regions, respectively. With further narrowing down the sequences containing Bra12 regulator binding sites using competition EMSA (Fig. 4BC and 4EF) and subsequently *in silico* searches using MEME, we were able to identify the 15-nucleotide Bra12 binding sequence, hereafter called the Bra12 box (Fig. 4G, right panel). All the identified boxes are located upstream of the previously designated TSSs (Fig. S11) in those gene promoters (Fig. S8 and Fig. 7), suggesting that the protein acts as positive transcriptional regulator for the corresponding genes. The two Bra12 boxes located in *bra12p* and the left portion of *bra0-1p* share identical sequences, while the third Bra12 box, located in the right portion of the *bra0-1p* region, differs in three positions from the former sites (Fig. 4G). Intriguingly, in the competition shift assay employing the radiolabeled DNA fragments comprising *bra12p* and *bra0-1p* regions (Fig. 4BE), we did not observe disruption of the corresponding protein-DNA complexes in the presence of respective bra12p_6-16 and bra0-1p_13-4 ctDNA fragments, containing the sequences overlapping the corresponding Bra12 boxes in those regions. However, we observed full out-competition of the Bra12 protein binding to those radiolabeled fragments when using longer ctDNA fragments (e.g., bra12p_6-15, bra0-1p_13-2) which were extended by a few or several nucleotides beyond the Bra12 boxes. These findings suggest an inefficient binding of the Bra12 protein to the sites located close to the ends of linear DNA fragments, and the need for the presence of flanking nucleotides necessary to stabilize those protein-DNA interactions.

**Fig. 4.**
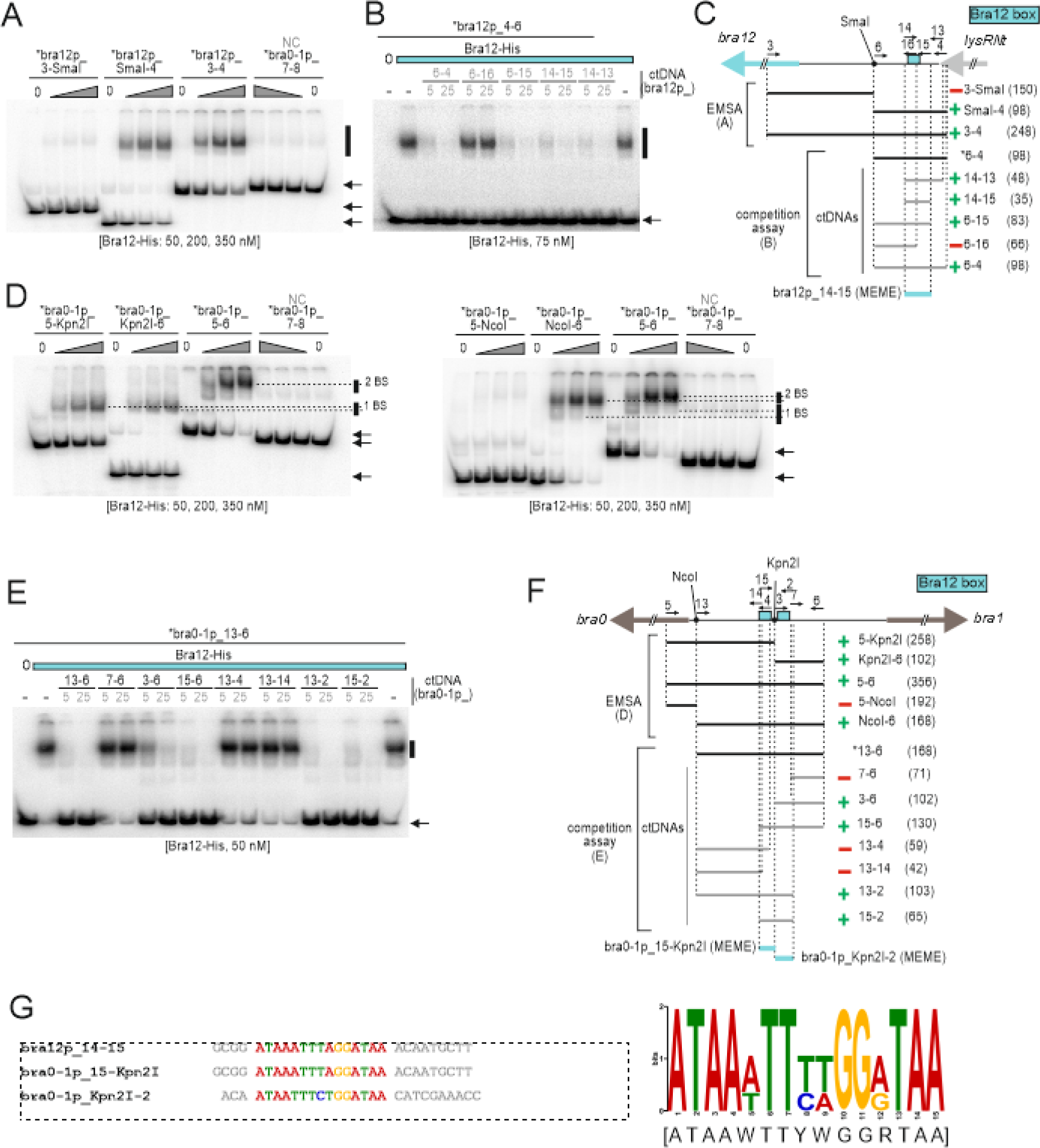
Determination of the Bra12 binding sequence. **(A-C)** and **(D-F)** present the identification of Bra12 binding sites within *bra12* and *bra0-1* promoter regions, respectively. **(A)** and **(D**) EMSA. The Bra12-His protein (concentrations shown below the gels) was incubated with the sets of *bra12p* or *bra0-1p* regions sub-fragments or a negative control (NC) fragment (bra0-1p_7-8, see also Fig. 1_2B). “1 BS” and “2 BS” in (D) refer to protein-DNA complexes formed by the DNA comprising 1 and 2 Bra12 binding sites, respectively. **(B)** and **(E)** competition EMSA. Bra12-His (75 nM, blue bar) was incubated with radioactively labeled bra12p_4-6 or bra0-1p_13-6 fragments (∼ 0.2 nM) and two sets of ‘cold-target DNA’ competitors (ctDNA) (5 or 25 nM) (PCR-amplified DNA indicated in gray font and lines). Black vertical bars and black arrows (A-B and D-E) indicate protein-DNA complexes and unbound DNA, respectively. **(C)** and **(F)** represent the summaries of panels A-B and D-E, respectively. Radiolabeled primers and DNA fragments are indicated with asterisks (*). The green symbols ‘**+**’ indicate Bra12-His interactions of Bra12-His with corresponding DNA fragments (EMSA) or the ability of the corresponding ‘cold-target DNA’ fragment to outperform the radiolabeled probe used (competition EMSA); the red symbols ‘**–**’ represent either the lack of protein-DNA interaction (EMSA) or the lack of the ability of the ‘cold-target DNA’ fragment to outperform the ^32^P-labeled DNA fragment (competition EMSA). The numbers next to the bars represent the primers used in the PCR reactions (Tab. S2); numbers in brackets represent fragments sizes (bp). **(G)** (left) Alignment of sequences containing the predicted Bra12 boxes. The MEME *in silico* tool was used to predict Bra12 sequences within EMSA-selected DNA fragments (fragments marked with blue bars in panels C and F). (right) Bra12 box sequence logo and the consensus sequence (below in square brackets) obtained using MEME.

In addition, due to the presence of non-overlapping Bra12 and SdpR boxes within the intergenic *bra0-1p* region, we also attempted to analyze the possibility of simultaneous binding of these proteins to the intergenic region. The EMSA experiments with a subfragment of *bra0-1p* (bra0-1p_5-6) comprising both Bra12 boxes and a proximally located single SdpR box revealed simultaneous and non-competitive binding of Bra12 and SdpR proteins to *bra0-1p* (Fig. S10A), and confirmed the relative locations of the Bra12 and SdpR boxes within that intergenic region (Fig. S10B).

Concluding, the above mentioned *in vitro* experiments allowed to determine the SdpR and Bra12 binding sites within *sdpR*, *bra0-1*, and *bra12* gene promoter regions. Given their locations relative to the gene TSSs in those promoter regions (Fig. 7), positive and complex regulatory functions in the transcriptional control for Bra12 and SdpR regulators have been proposed.

#### Impact of SdpR and Bra12 on gene promoter *activity*

Protein-DNA interaction analyses allowed a precise localization of the SdpR and Bra12 regulator binding sites within the promoters of the genes crucial for brasilicardin biosynthesis (*bra0*, *bra1,* and *bra12*). To investigate their impacts on target gene promoters, we applied bacterial luciferase reporter assay. The well-established and easily genetically tractable *S. coelicolor* strain M1154 was used. The strains comprised either the wild-type bcaAB01 fosmid or gene deletion versions of that fosmid (bcaAB01_*ΔsdpR* and bcaAB01_*Δbra12*) together with the appropriate reporter pFLUXH plasmids containing luciferase encoding gene operon *luxCDABE* (Craney *et al*., 2007) under the control of a studied gene promoter.

The time-course analyses for the reporter strains showed a negative impact of the *bra12* gene deletion on the activities of the *bra0* and *bra1* gene promoters, as well as its own gene promoter, *bra12p* (Fig. 5A). The deletion of *bra12* most significantly affected *bra1p* for which up to a 100-fold reduction in luminescence levels was observed in comparison to the wild-type strain (Fig. 5B). A less pronounced effect of *bra12* gene deletion was observed in the case of the *bra0p* and *bra12p* (up to a 10-fold decrease in both cases) (Fig. 5AB). The negative impact on gene promoter activities was also observed in the case of *sdpR* gene deletion strains comprising *bra0p* or *bra12p* (Fig. 5A), manifested by 10- and 100-fold reductions, respectively (Fig. 5B). The essential role of SdpR for *bra12p* activity was also observed in the reporter strain comprising the *sdpR* gene under the constitutive promoter *ermE*\* (Fig. S9). Interestingly, while only minor differences in luminescence levels of the *bra1p* region between *ΔsdpR* and WT strains were observed, the deletion of *sdpR* exerted a small-time shift in the luminescence activity peak (Fig. 5A, middle panel).

**Fig. 5.**
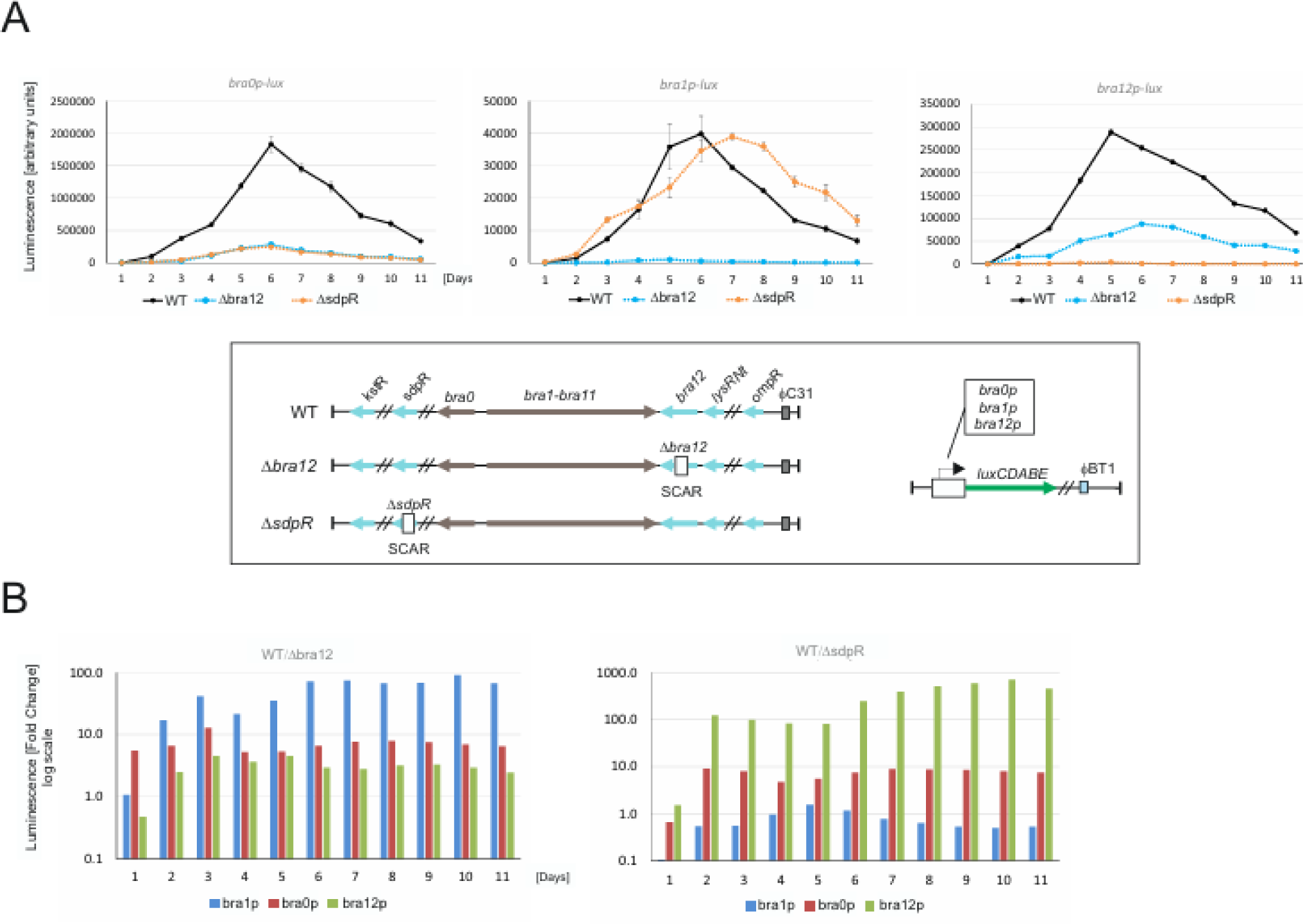
Impact of SdpR and Bra12 regulators on the activities of gene promoters in the Bra-BGC. Luciferase assays. **(A)** Measurements were conducted using heterologous strains of *S. coelicolor* M1154 that harbor either a *sdpR* or a *bra12* gene deletion fosmid (bcaAB01_ΔsdpR or bcaAB01_Δbra12, respectively), and the corresponding gene promoters (*bra0p*, *bra1p* and *bra12p*) delivered onto pFLUXH integrative plasmids. The luminescence readings were normalised against the OD_600_ of the corresponding cultures grown on solid DNA medium for 11 days. The frame at the bottom depicts the fosmids and plasmids used to create corresponding reporter strains. **(B)** The fold changes of promoter activities. The graphs were created using averaged luminescence values obtained for corresponding time points.

In sum, the gene expression analyses using reporter strains indicated the impacts of both Bra12 and SdpR regulators on the activities of the promoters controlling the genes essential for brasilicardin biosynthesis. The absence of the Bra12 regulator exerts a negative impact on the activities of *bra0*, *bra1*, and *bra12* gene promoters, suggesting that Bra12 acts as a positive transcriptional regulator. The absence of SdpR, negatively impacts the activities of the *bra0* and *bra12* gene promoters and exerts a negligible impact on the *bra1* promoter, indicating that the protein presumably plays a complex overall role in gene expression.

### 3.3 SdpR and Bra12 impact brasilicardin production

To study the influence of regulatory proteins on biosynthesis, we evaluated the production levels of brasilicardin congeners in liquid cultures of several mutant strains. Due to the low genetic tractability of the *N. terpenica* strain, we used the previously established heterologous host *Amycolatopsis japonicum*, also belonging to Actinomycetota (Schwarz *et al*., 2018c; Schwarz *et al*., 2018a). To examine the impact of a lack of regulators, we individually introduced to the host strain the bcaAB01 fosmids comprising *sdpR* or *bra12* gene deletions (bcaAB01_*ΔsdpR* and bcaAB01_*Δbra12*) (Tab. S1). To study overexpression of the regulatory genes, we cloned the corresponding genes into the integrative plasmid pIJ10257 under the constitutive *ermE*\* promoter (*ermE*\*p) (pIJ_bra12, and pIJ_sdpR), and introduced the obtained “overexpression” plasmids into the *A. japonicum* host strain carrying the pPS1 derivative of bcaAB01 fosmid (Tab. S1). The pPS1 facilitates analyses of the impact of individual regulators as it comprises only a minimal set of genes essential for brasilicardins biosynthesis (*bra0*-*bra12*) and confers production of brasilicardins on levels similar to those of the strain harboring the bcaAB01 fosmid (Schwarz *et al*., 2018a). To analyze the effects of gene deletions, the corresponding derivatives of the bcaAB01 fosmid were individually transferred to the wild-type *A. japonicum* host.

HPLC/MS analysis of liquid culture extracts of the *bra12* overexpression strain (*A. japonicum*::pPS1+pIJ_*bra12*) revealed significantly higher (296%) levels of brasilicardins compared to the control strain harboring solely the empty plasmid pIJ10257 (*A. japonicum*:: pPS1 + pIJ) (100%) (Fig. 6). In support of this result, the *bra12* deletion mutant created in our lab (*A. japonicum*::bcaAB01_*Δbra12*) showed no production of brasilicardins, which is also in line with the previous findings (Schwarz *et al*., 2018a). Conversely, the SdpR overexpression strain (*A. japonicum*::pPS1+pIJ_*sdpR*) exhibited a negative effect manifested by a clear drop (64%) in brasilicardin production, while the deletion mutant of this gene (*A. japonicum*::bcaAB01_*ΔsdpR*) showed pronouncedly higher brasilicardin levels (397%) with the reference to the control strain.

**Fig. 6.**
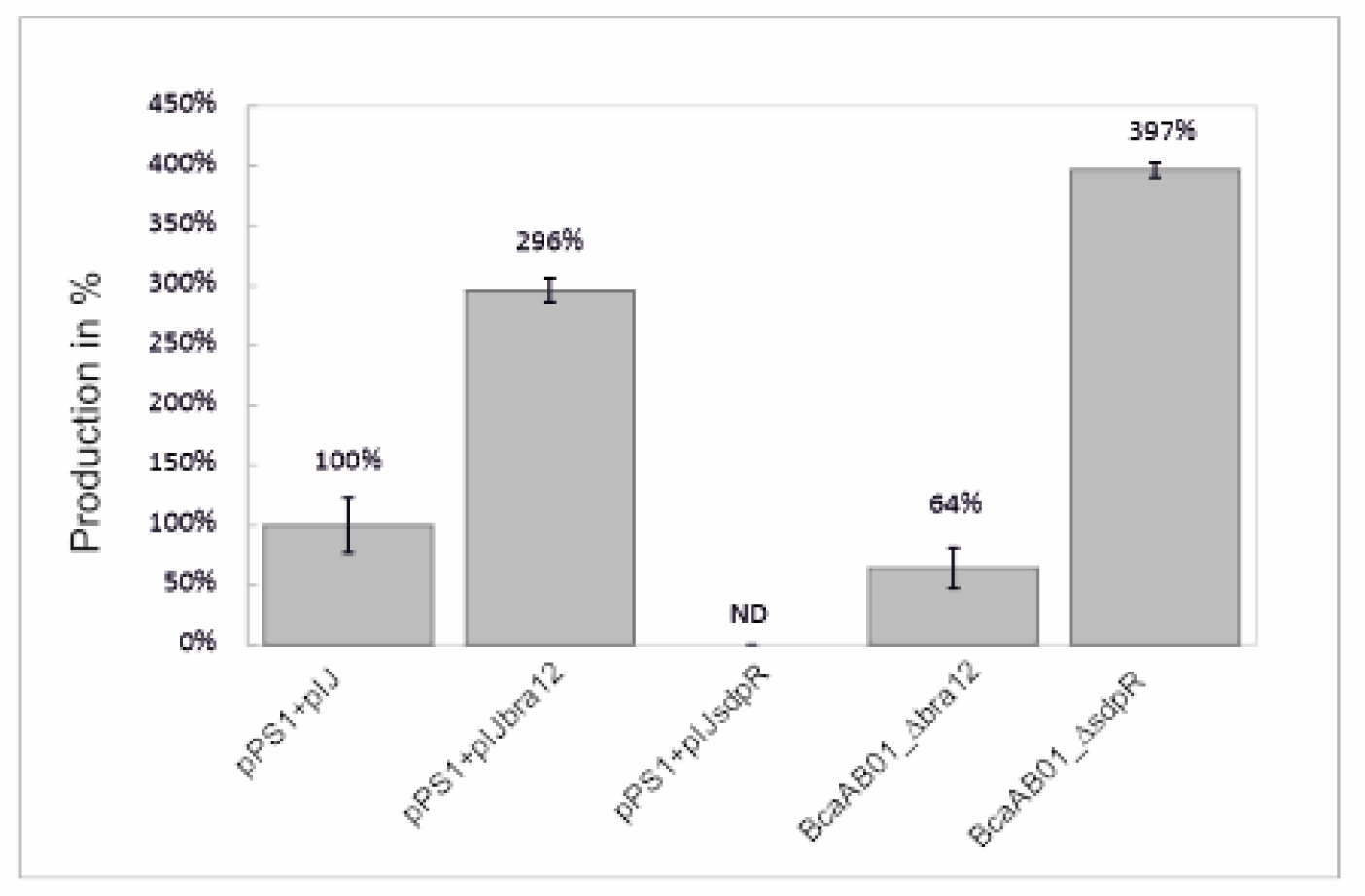
The influence of regulatory proteins Bra12 and SdpR on the biosynthesis of brasilicardin congeners. HPLC/MS. Total production of brasilicardins (sum of BraC, Brac-agl, BraD, BraD-agl) was measured in the supernatants of 72-h liquid cultures of *A. japonicum* strains harboring corresponding constructs (see Tab. S1). The bars represent the averages of three technical replicates obtained for each strain and normalised to *A. japonicum*::pPS1+pIJ (100%), the standard deviations were calculated using Excel. ND is non-detectable.

To sum up, we confirmed the essential role of the Bra12 protein for brasilicardin biosynthesis using a heterologous host expression system. We also showed that the SdpR regulator negatively affects the production of the corresponding secondary metabolites.

## 4. Discussion

Brasilicardin A is considered a promising immunosuppressive lead structure for organ transplant therapies. The recent findings showed a semisynthetic approach as the effective way to obtain a complete compound in laboratory scale quantities (Botas *et al*., 2021). However, further improvements regarding the production yields are limited by the availability of biosynthetically generated brasilicardin intermediates. Amelioration in their production levels seems to be hampered to the greatest extent by the lack of understanding of molecular mechanisms regulating gene expression in the brasilicardin gene cluster, Bra-BGC. The sequence analysis of the gene cluster and its flanking regions revealed the presence of multiple genes encoding putative transcriptional regulators (KstR, SdpR, Bra12, LysRNt and OmpR) suggesting a complex transcriptional regulatory network (Wolański *et al*., 2021). However, only the roles of the LysRNt and Bra12 have been investigated to date, in addition, to a various extent. While, the role of LysRNt regulator in brasilicardin biosynthesis, and its presumed mechanism controlling activity of the protein has been revealed in a recent study (Wolański *et al*., 2021), the function of Bra12, considered as the major positive regulator, has not been explored in detail (Schwarz *et al*., 2018a). To broaden our understanding of the regulatory dependencies in the Bra-BGC, we embarked on the investigation of the unstudied putative regulators KstR, SdpR, OmpR, and delved into the function of Bra12.

To elucidate the potentials of new regulatory candidates and Bra12, to control gene expression within the Bra-BGC, we first looked for the binding sites for those proteins within the gene cluster. Using EMSA approach, we demonstrated that KstR, SdpR and Bra12 proteins, were able to interact with several gene promoters located within the regions comprising the Bra-BGC and flanking genes on the fosmid BcaAB01. Nevertheless, we did not detect binding of OmpR protein to any of the fosmid regions. This led to subsequent exclusion of this putative regulator from further analyses. Noteworthy, the deletion of the BcaAB01 region comprising the *ompR* gene did not affect brasilicardin biosynthesis in the previous study (Schwarz *et al*., 2018a), suggesting that the OmpR does not play function in the compound production. The second regulator, KstR of TetR family, was also excluded from further investigation, as it only exhibited potential to regulate expression of a putative gene operon *33470*-*33475-kstR*, including its own gene. However, the glutamine-dependent synthetase and peptidyl-arginine deaminase enzymes, encoded by *33470*-*33475*, usually play roles in nitrogen (amino-acid) metabolism pathways, and according to a previous study are not related to precursor supply and brasilicardin biosynthesis (Schwarz *et al*., 2018b). The two other candidate regulators, Bra12 and SdpR, showed greater potential to regulate gene expression in the Bra-BGC as they exhibited binding to the promoters of biosynthetically essential genes *bra0-bra1* and *bra1,* and their own gene promoter regions, suggesting in addition possible transcriptional autoregulation. Moreover, SdpR exhibited binding to *bra12* promoter region indicating another level of transcriptional regulation in the gene cluster.

An initial prediction of Bra12 and SdpR functions, based on the locations of their binding sites in relation to the transcriptional start sites (TSS) determined using RNA-seq, led us to propose that Bra12 plays positive, while SdpR not clearly defined role in gene expression. Indeed, quantitative analysis using reporter gene expression system, showed that the strains lacking *bra12* gene exhibit pronouncedly decreased activity of *bra0*, *bra1* and *bra12* gene promoters (∼10, ∼100 and ∼10-fold, respectively), in comparison to the control strains containing the gene. The results correspond to a previous qualitative gene expression study that showed a crucial role of Bra12 in the transcription of the entire set of genes involved in the brasilicardin biosynthesis, namely *bra0* and *bra1-bra11*, and its own *bra12* gene (Schwarz *et al*., 2018a). We assume, that the strongest effect observed for *bra1* promoter emphasizes the essential role of Bra12 not only in the expression of gene *bra1*, but also for the entire *bra1-bra11* biosynthetic gene cluster, as those genes are assumed to form operon or sub-operon (Wolański *et al*., 2021). Disruption of *bra12* affects negatively also, though less pronounced (∼10-fold change), the activity of the promoter of another gene, this time *bra0*, orientated divergently to *bra1*, and sharing within *bra1* a common intergenic region. Noteworthy, *bra0* codes for the dioxygenase involved in decorating brasilicardin congeners with the methoxy group at carbon atom C16. Although, Bra0 does not play an essential role in brasilicardin backbone biosynthesis (Schwarz *et al*., 2018a), the methoxy group seems to be crucial for the immunosuppressive activity of brasilicardin A (Komatsu *et al*., 2004; Komatsu *et al*., 2005). Altogether, our findings indicate that Bra12 controls the activity of all biosynthetic genes in the gene cluster. We propose, that binding of the Bra12 within the *bra0-1* region is required to activate expression of the entire Bra-BGC and suggest that Bra12 alone might exert a major impact on brasilicardin biosynthesis. In addition, the activity of the *bra12* gene promoter shows a positive correlation with the presence of Bra12, indicating autoregulation of its own gene (Fig. 5). The requirement of Bra12 for the transcription of *bra12* gene was first observed in a previous study (Schwarz *et al*., 2018a). We demonstrate that this is likely due to a direct regulation, as Bra12 binds to the *bra12* gene promoter region. Transcriptional autoregulation was reported for several regulators of SARP (Streptomyces Antibiotic Regulatory Protein) family to which Bra12 belongs to (Santamarta *et al*., 2002; Hoskisson *et al*., 2006; Novakova *et al*., 2010; Tsypik *et al*., 2021). It’s also worth mentioning, that some of the known AfsR/SARP regulators may undergo phosphorylation, required to enhance their DNA-binding activity. The analysis using the SMART tool identifies a N-terminal DNA binding domain of Bra12 as Trans_reg_C receiver domain (Pfam PF00486), typical for response regulators of two component phosphorelay transduction systems. In addition, our sequence analysis has identified at least 21 homologs of Bra12 encoded in *N. terpenica* genome. This suggests that AfsR homologs can likely form a number of regulatory networks responding to various environmental stimuli in this organism. However, putative activation of Bra12 by phosphorylation, and further impact on expression of the Bra-BGC has not been investigated and remains unknown. Importantly, heterologous production assays confirm essential role of Bra12 for the biosynthesis of brasilicardins (see further discussion).

In the case of SdpR, the arrangement of its binding sites in the promoter regions did not allow to clearly predict a general function for the protein in the regulation of gene expression in the Bra-BGC. For instance, in the biosynthetically essential *bra0-1* region, the presence of two SdpR boxes, both upstream and downstream *bra0* and *bra1* gene TSSs, respectively, suggested a corresponding positive and negative role in gene expression (Fig. 7). However, the reporter assays revealed a negative impact of *sdpR* gene disruption on the activity of the *bra0* promoter (∼10-fold decrease in comparison to control strain), while for *bra1* only a slight delay in the promoter activity peak was observed (Fig. 5A). The impact exerted on the *bra0* promoter corresponds to the predicted function for the SdpR. We propose that two effects presumably exerted the observed shift in the *bra1* promoter activity, a reduction of the level of Bra12 and an absence of SdpR. The reduction of Bra12 levels might result from a pronouncedly decreased (over 100-fold) activity of the *bra12* promoter observed in *ΔsdpR* strain. Importantly, the positive impact of SdpR on *bra12p* activity was also confirmed using a *sdpR* overexpression strain (Fig. S9). Further, a decreased level of Bra12 in the absence of SdpR should therefore lead to a significant reduction in *bra1* promoter activity. However, we demonstrate that lack of SdpR leads only to a small delay in *bra1p* activity, and finally to increased levels of brasilicardins. These findings led us to suggest that SdpR may negatively, by binding to the SdpR box located downstream the TSS of the *bra1* gene, regulate expression of this gene promoter. In sum, the combined impact of the absence of SdpR repressor and decrease in Bra12 activator levels results in a slightly delayed activation of the *bra1* promoter. Importantly, using EMSA we demonstrate that both regulators does not compete for binding to DNA fragments comprising the *bra0-1* intergenic region (Fig. S10). This suggests that Bra12 and SdpR may independently exert impact on gene expression of *bra1* (and the downstream genes *bra2*-*bra11*) probably as the result of a balance in cellular levels between both proteins. The complexity of putative regulatory functions by SdpR has been also revealed in the case of *sdpRp*. Surprisingly, despite the presence of two SdpR boxes in the *sdpRp* region, SdpR did not show any impact on its own gene promoter activity excluding the possibility of an autoregulatory function. We speculate that SdpR, that belongs to a large family of metalloregulators wide-spread in bacteria, may, by subject to another level of regulation, be related with the metal sensing as it was shown for ArsR, SmtB and CmtT regulators of *E. coli*, *Synechococcus*, and *Mycobacterium*, respectively (Erbe *et al*., 1995; Xu *et al*., 1996; Wang *et al*., 2005).

**Fig. 7.**
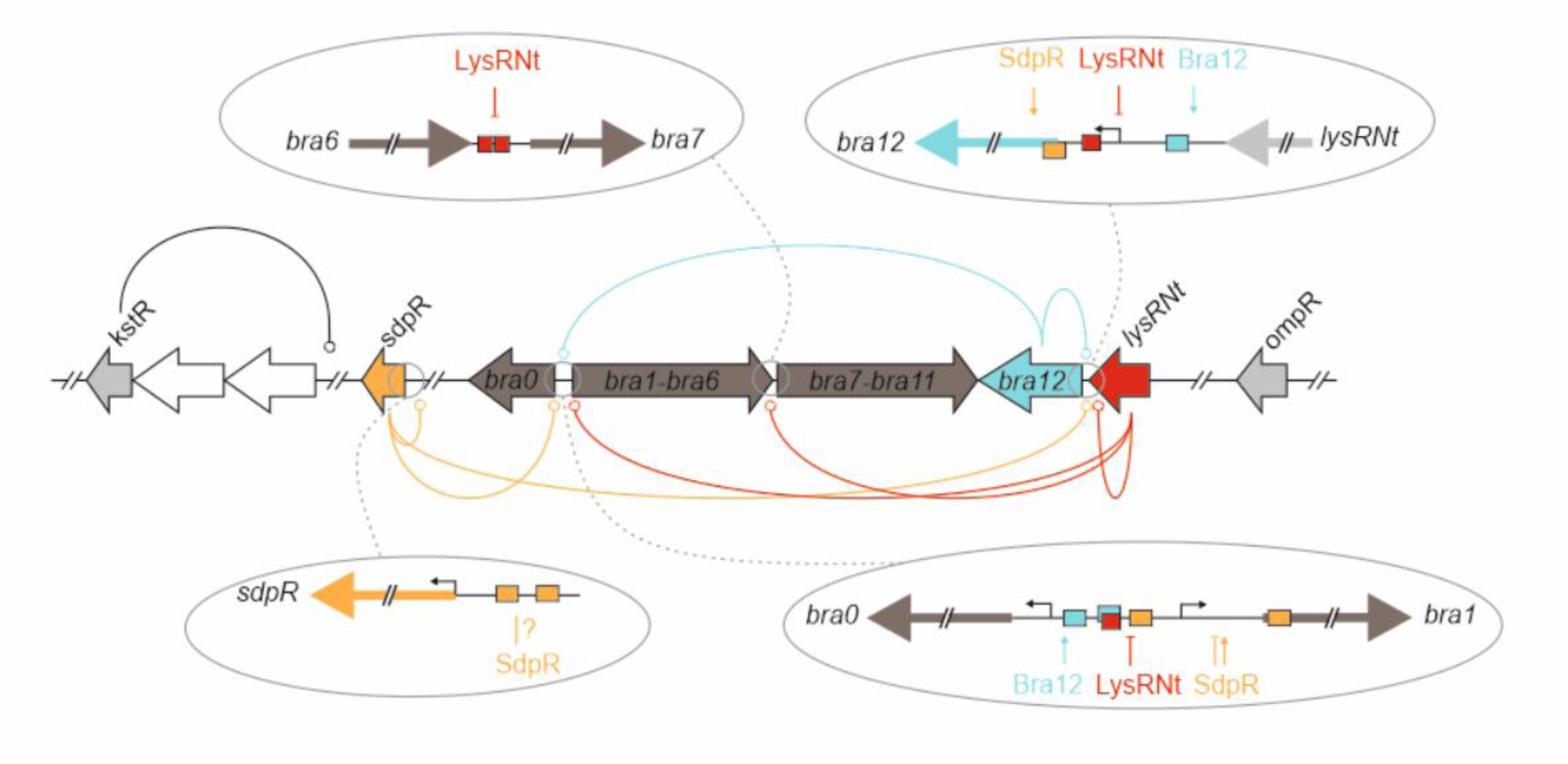
Presumable regulation of the Bra-BGC. The genes encoding regulatory proteins are linked to the identified target gene promoter regions using colour-coded solid lines that end up with circles. The blue, orange and red rectangles within the promoter regions depict the locations of the binding sites of the regulatory proteins Bra12, SdpR and LysRNt, respectively. The assumed or verified effects exerted by the regulators on activities of the corresponding gene promoters are presented by vertical arrows (activation) and T-lines (repression). In the case of LysRNt, its putative role in gene transcription has been shown, as the direct impact on promoter activity was not studied (Wolański *et al*., 2021). The bent arrows indicate the transcription start sites (TSS) (see also Fig. S8).

Here, we demonstrate, that both Bra12 and SdpR exert important, nevertheless opposite effects, on brasilicardin congener biosynthesis. Consistently, with the previous research (Schwarz *et al*., 2018a), we show that Bra12 positively affects the process of biosynthesis, although, in comparison to that study, we observed about a 3-times higher increase in the production titers in the host strain overproducing the Bra12 regulator (63 vs 196%, respectively) (Fig. 6). As in both studies, the *bra12* was delivered under constitutive *ermE*\* promoter on the pIJ10257 integrative plasmid, the discrepancies in biosynthesis yields likely result from various genetic backgrounds of the host strains. Indeed, a full length wild-type bcaAB01 fosmid (Bra-BGC plus flanking genes) was used in the previous study, while a shortened pPS1 version (only the Bra-BGC) was employed in our approach. This clearly indicates additional impact exerted by other regulatory proteins encoded in the genes flanking the Bra-BGC. Up to date, the LysRNt, and SdpR are the only, apart from Bra12, identified regulators of the brasilicardin gene cluster. Importantly, individual overexpression of LysRNt and SdpR regulators in the identical genetic backgrounds (host strain harboring pPS1) showed their negative impacts manifested by a decrease in the brasilicardin production levels (∼27% and 64% levels of the strain harboring only the pPS1 fosmid for LysRNt and SdpR overexpression, respectively). Consistently, deletion of SdpR leads to a pronounced increase in the production levels of brasilicardin congeners (∼397%).

<u>Final conclusions</u>. Regulation of gene expression in the Bra-BGC seems complex, and represents a still not fully understood process that requires concerted action of multiple regulators to ensure biosynthesis of the BraA compound. We assume that the intergenic *bra0-1* region plays a role of an essential regulatory “hot-spot” comprising binding sites for multiple regulatory proteins that control transcription of the biosynthetic genes in a coordinated way. The involvement of multiple regulator binding sites within this region may reflect the complexity of conditions of this free-living bacterium which it faces in the environment and the necessity of diversified responses to external factors to ensure tight control over gene expression in the Bra-BGC. As the putative function of the BraA during infection has not been explained so far, the range of external signals may also include the ones related with the host organism of this facultative pathogen. In addition, the feedback mechanisms including regulation of gene expression of regulatory genes by other regulators creates another level of complexity and makes understanding of transcriptional regulatory circuits taking place in Bra-BGC even more challenging. To contribute to this process, we propose a current model for the network of regulatory dependencies in the Bra-BGC (Fig. 7).

## Acknowledgement

We thank to Constantin Kiel (University of Bielefeld) for purification of Bra12-His and initial EMSA tests. The work from Harald Gross were funded by Bundesministerium für Bildung und Forschung (BMBF) (FKZ 031A568A), while results from Marcin Wolański and Jolanta Zakrzewska-Czerwińska were funded by National Centre for Research and Development (NCBR) (ERA-NET-IB/NeBrasCa/10/2015).

## 5. Conflict of interest

The authors declare that they have no competing interests.

## Supplementary figures

**Fig. S1.**
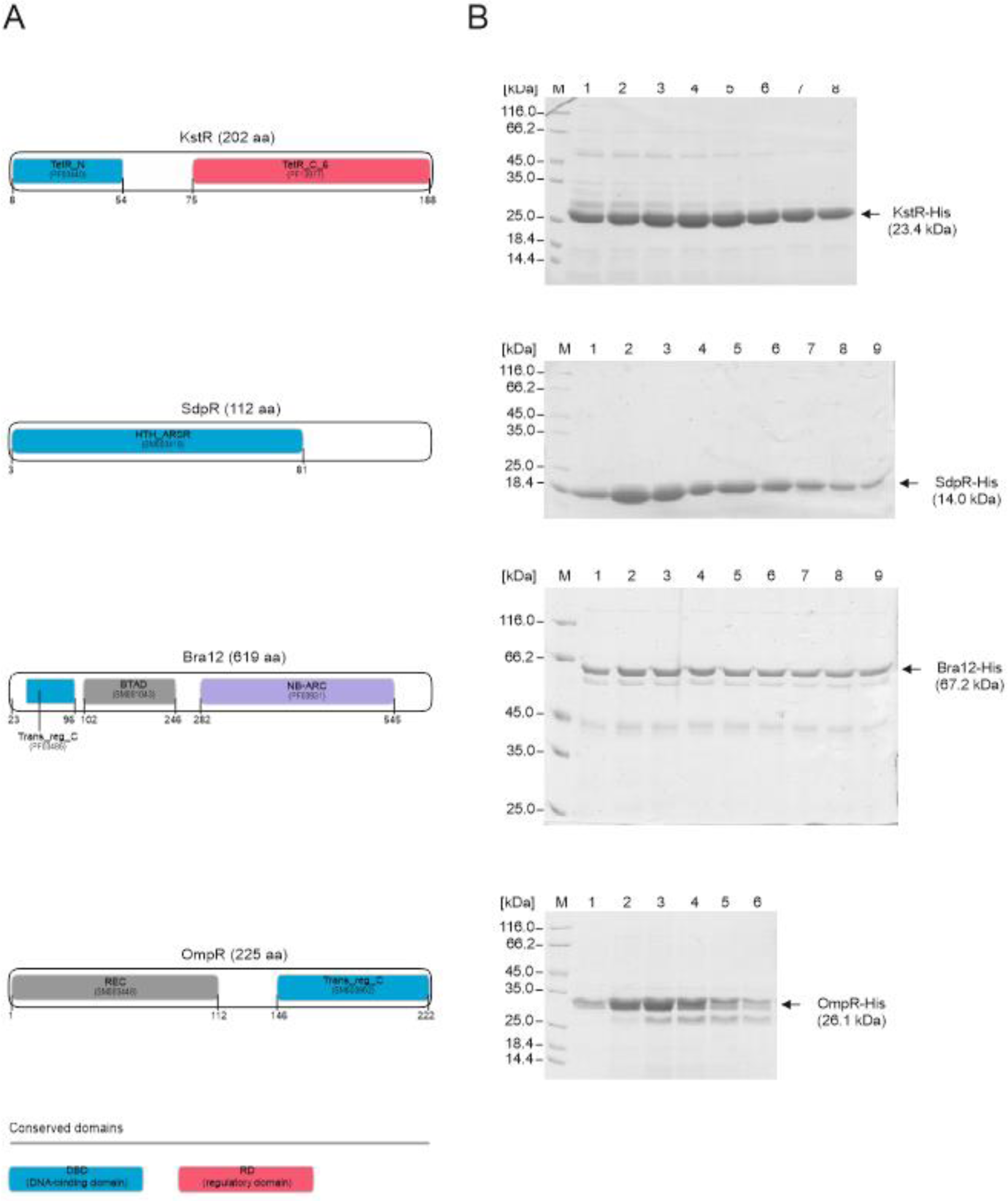
Purification of recombinant proteins. **(A)** Schematic depiction of protein domain organizations based on SMART and conserved domain (CD) searches (see SI). The SMART and Protein Family (PF) numbers of the identified protein domains are given. The DNA-binding domain has been indicated with blue and other domains according to the legend. **(B)** SDS-PAGE analysis of purified recombinant proteins. The His-tagged proteins were purified using metal affinity resins, HiTrap Talon® crude column (1 ml) or His-Select® Nickel Affinity Gel, as described in detail in SI. The elution fractions were collected while the resins were washed with an increasing gradient of buffer B (2-50%) containing 500 mM imidazole. The arrows indicate bands representing the corresponding recombinant proteins together with their molecular weights. M: protein weight marker (#26610, Thermo Fisher Scientific).

**Fig. S2.**
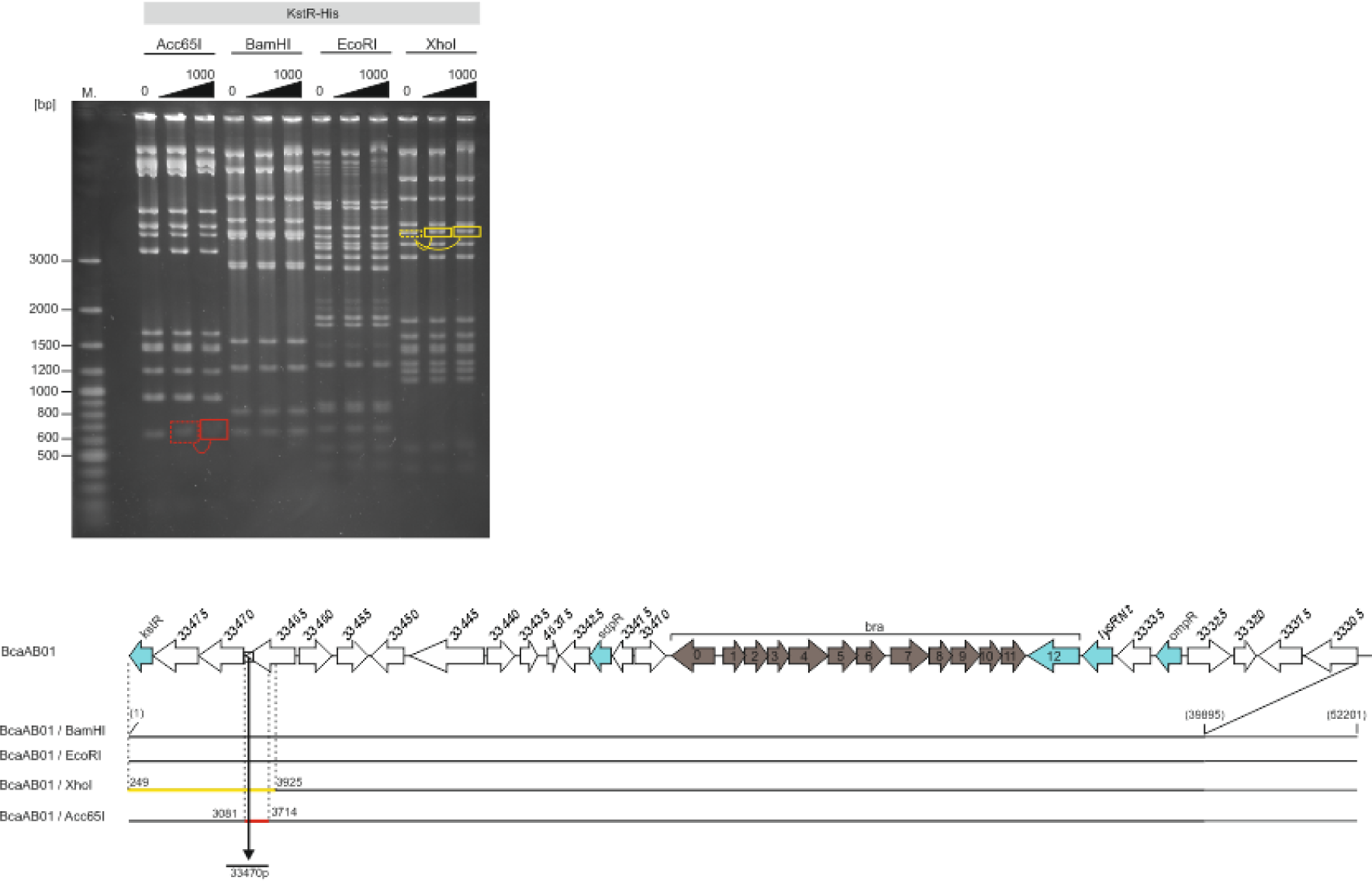
Preliminary identification of the KstR protein binding sites within the bcaAB01 fosmid. EMSA. The fosmid was digested independently with four restriction enzymes (Acc65I, BamHI, EcoRI, and XhoI) and incubated in the presence of 100 and 1000 nM concentrations of KstR-His protein, followed by electrophoresis on an agarose gel. The DNA was visualized by soaking the gel in an ethidium bromide solution. The red and yellow rectangles indicate vanished and shifted bands, respectively. M: DNA molecular weight marker (#SM0323, Thermo Fisher Scientific). The bottom panel represents a graphical representation of the brasilicardin biosynthetic gene cluster on bcaAB01 fosmid and identified vanished and shifted DNA fragments (red and yellow bars, respectively). The numbers next to those bars show the nucleotide positions on bcaAB01. Numbers above the genes are NCBI accession numbers in “AWN90_RS…” format. The gene promoters selected for further analysis are indicated by a black vertical arrow.

**Fig. S3.**
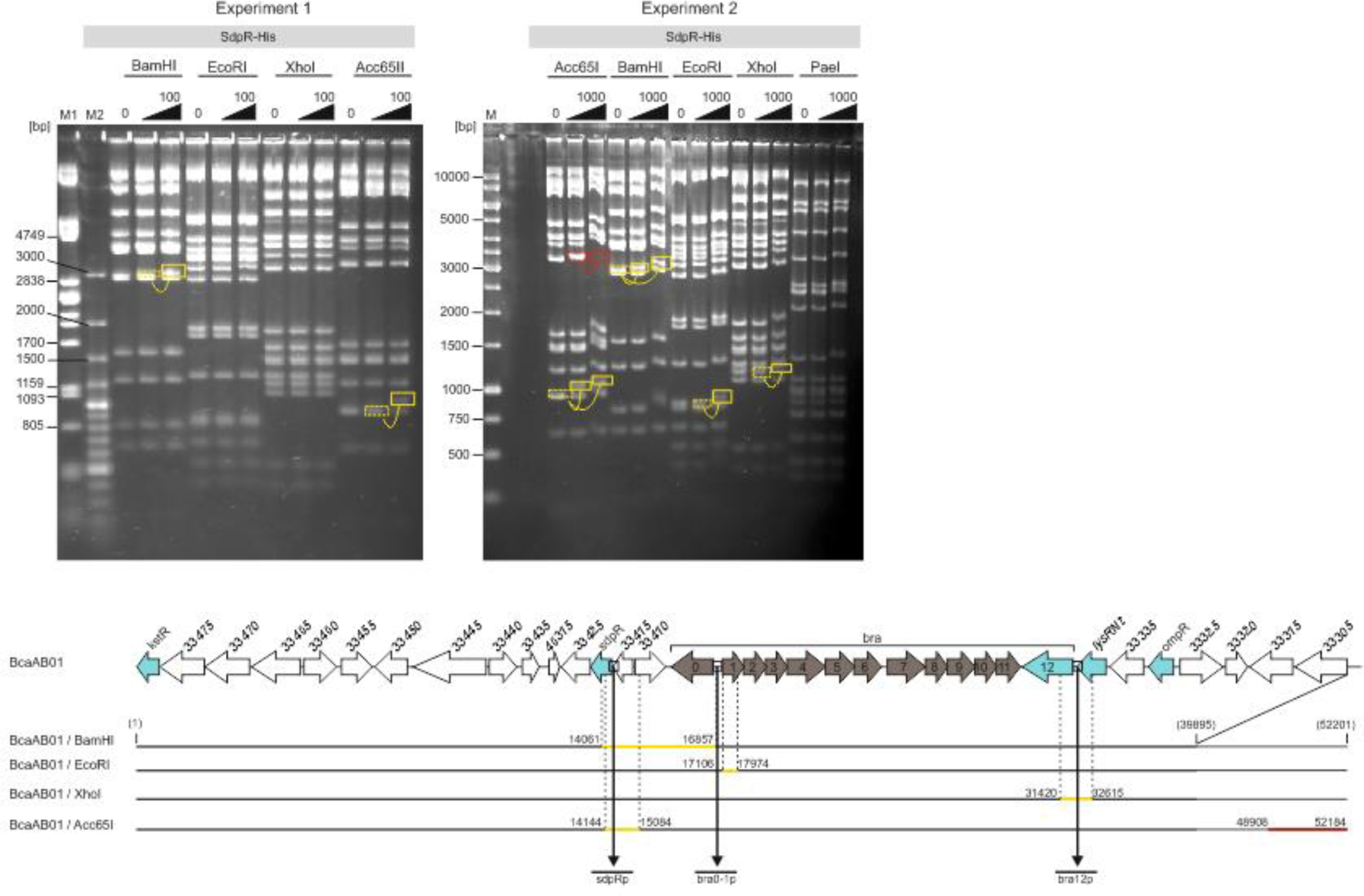
Preliminary identification of the SdpR protein binding sites within the bcaAB01 fosmid. Electrophoretic mobility shift assay. The figure represents two experiments. In each of those, the fosmid was independently digested with a set of different restriction enzymes and incubated in the presence of increasing concentrations of SdpR-His protein (10 and 100 nM – experiment 1; 100 and 1000 nM – experiment 2). Incubation was followed by electrophoresis on an agarose gel, and DNA visualisation was conducted by soaking the gel in ethidium bromide solution. The red and yellow rectangles on the gels indicate vanished and shifted bands, respectively. M1, M2 and M – DNA molecular weight markers (λ/PstI, # 3530-500, A&A Biotechnology; #SM0323, Thermo Fisher Scientific – experiment 1, and #SM0311, Thermo Fisher Scientific – experiment 2, respectively). The bottom panel represents a graphical analysis of the results. The numbers next to those bars show the nucleotide positions on bcaAB01. Numbers above the genes are NCBI accession numbers in “AWN90_RS…” format. The gene promoters selected for further analysis are indicated by a black vertical arrow.

**Fig. S4.**
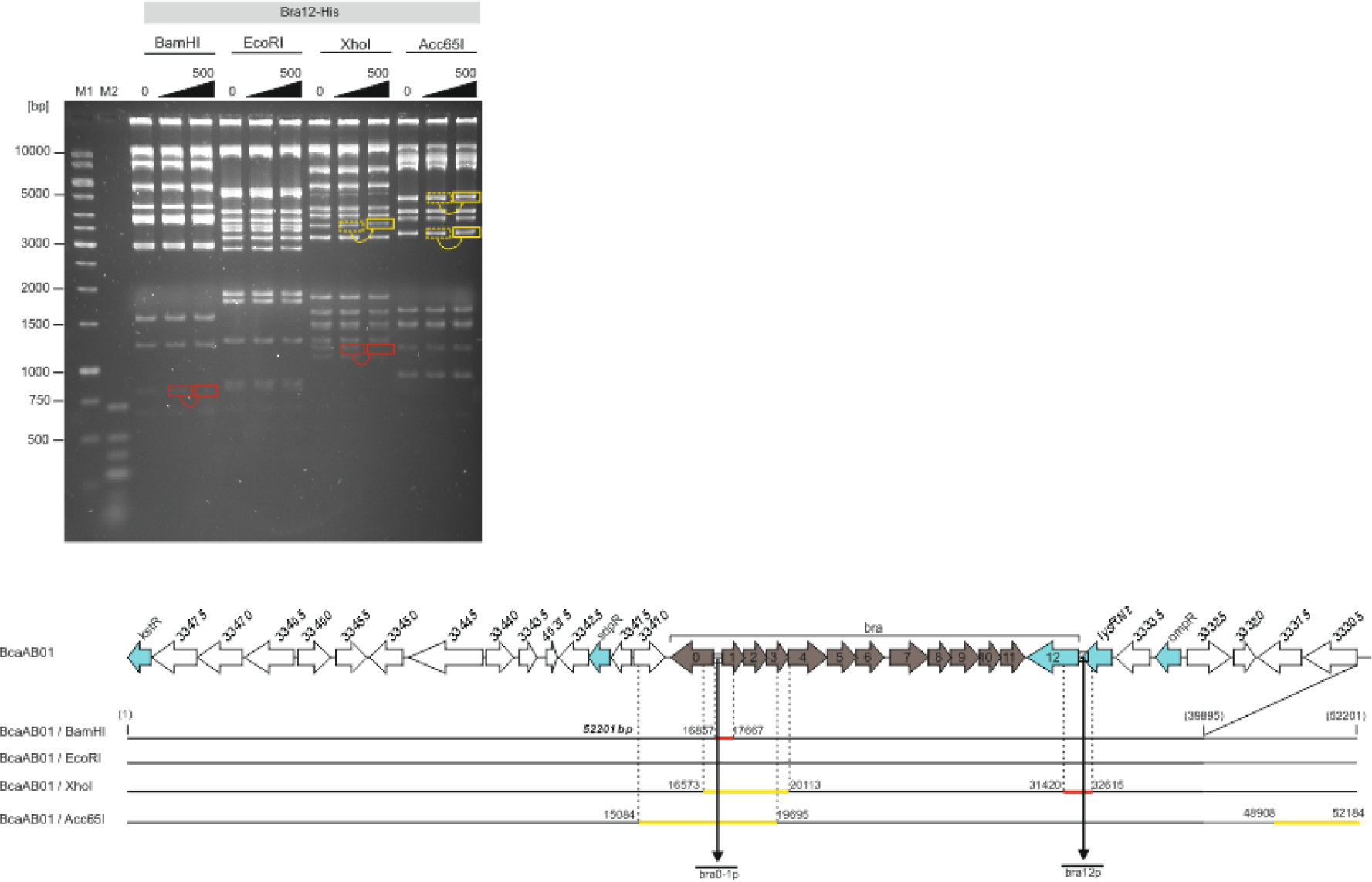
Preliminary identification of the Bra12 protein binding sites within the bcaAB01 fosmid. Electrophoretic mobility shift assay. The fosmid was independently digested with four restriction enzymes (Acc65I, BamHI, EcoRI, and XhoI) and incubated in the presence of 50 and 500 nM concentrations of Bra12-His protein, followed by electrophoresis in an agarose gel. The DNA was visualised by soaking the gel in an ethidium bromide solution. The red and yellow rectangles indicate vanished and shifted bands, respectively. M1, M2 – DNA molecular weight markers (#SM0311, Thermo Fisher Scientific; #SM1193, Thermo Fisher Scientific). The bottom panel represents a graphical analysis of the results. The numbers next to those bars show the nucleotide positions on bcaAB01. Numbers above the genes are NCBI accession numbers in “AWN90_RS…” format. The gene promoters selected for further analysis are indicated by a black vertical arrow.

**Fig. S5.**
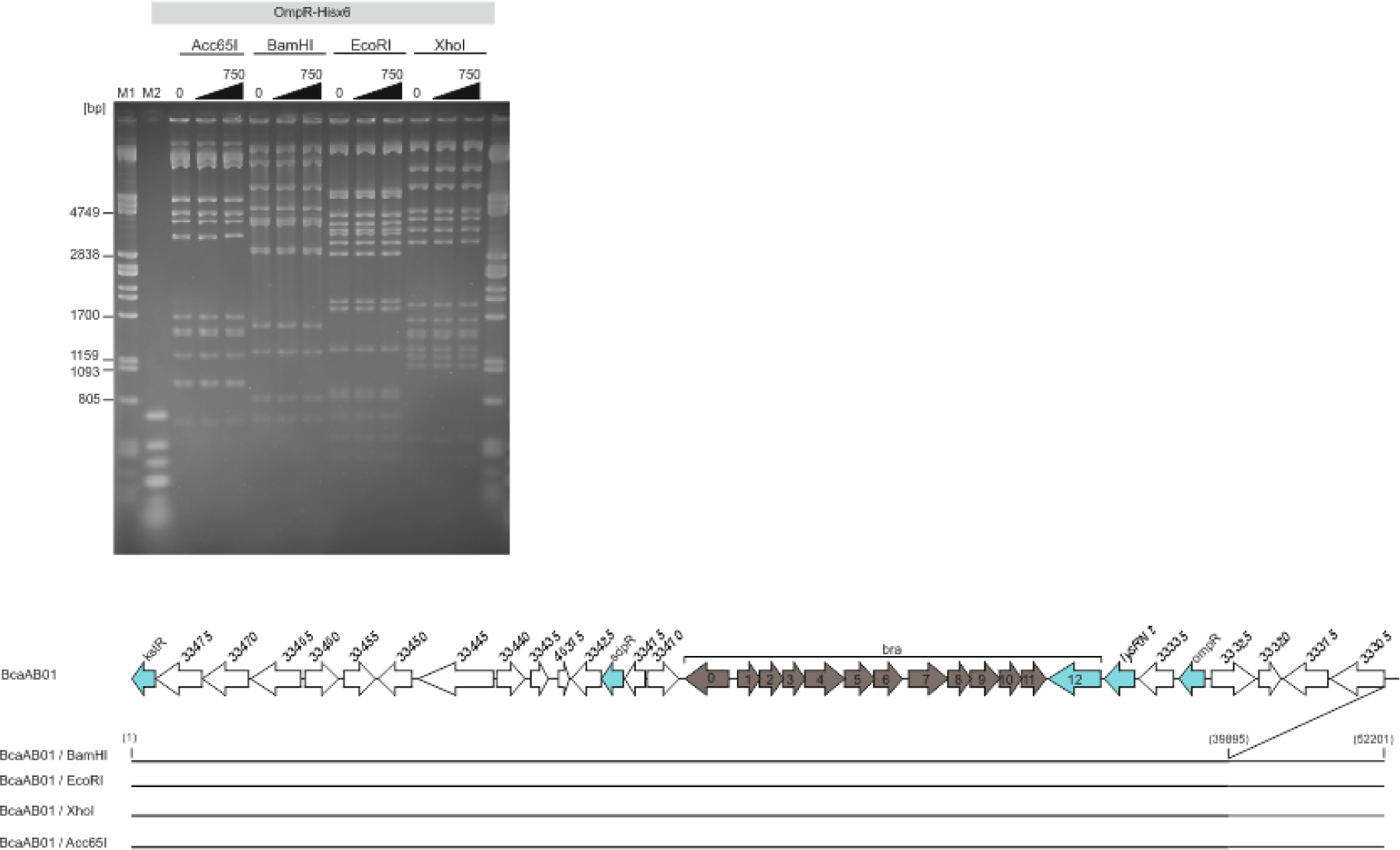
Preliminary identification of the OmpR protein binding sites within the bcaAB01 fosmid. Electrophoretic mobility shift assay. The fosmid was independently digested with four restriction enzymes (Acc65I, BamHI, EcoRI, and XhoI) and incubated in the presence of 100 and 750 nM concentrations of OmpR-His protein, followed by electrophoresis on an agarose gel. The DNA was visualised by soaking the gel in an ethidium bromide solution. The red and yellow rectangles indicate vanished and shifted bands, respectively. M1, M2: DNA molecular weight markers (λ/PstI, # 3530-500, A&A Biotechnology; #SM0311, Thermo Fisher Scientific; #SM1193, Thermo Fisher Scientific). The bottom panel represents a graphical analysis of the results. Numbers above the genes are NCBI accession numbers in “AWN90_RS…” format.

**Fig. S6.**
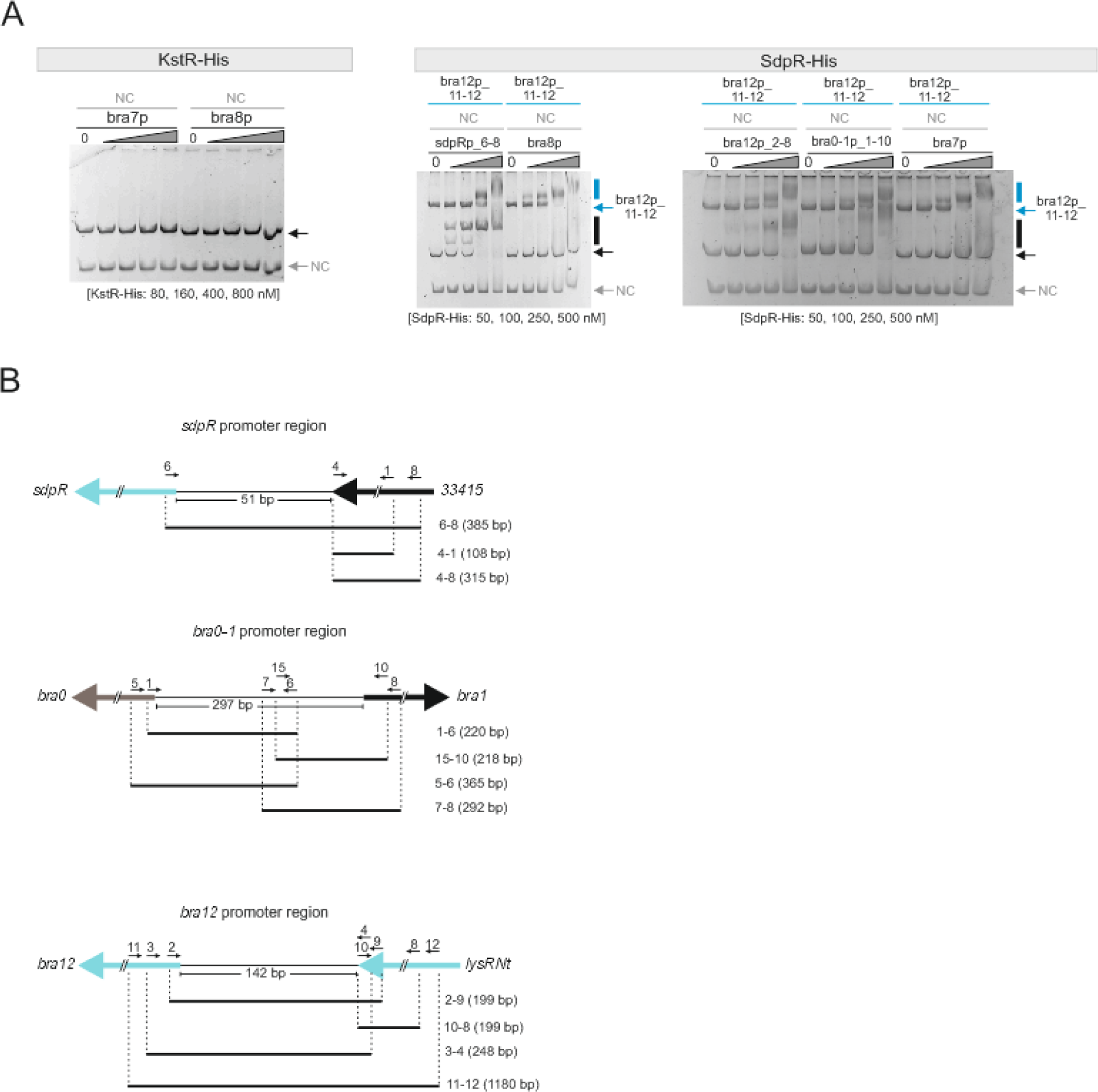
Supplementary identification of regulatory protein target promoters within the Bra-BGC. **(A)** Electrophoretic mobility shift assays (EMSA). Recombinant proteins (KstR-His and SdpR-His) were incubated with preselected promoter regions (see Fig. S2-5) amplified by PCR. In the assays, constant amounts of unlabeled DNA and varying concentrations of proteins were used, as indicated. To confirm specific binding and compare binding to different DNA fragments, the reaction mixtures were spiked additionally with the DNA comprising the *33140* gene promoter region (negative control, NC) (KstR and SdpR panels) and other DNA fragments comprising promoters of the Bra-BGC (only SdpR panel). Vertical black bars indicate protein-DNA complexes, and black and gray arrows indicate unbound DNA fragments. The DNA fragments and corresponding protein-DNA complexes are colour-coded. **(B)** Graphical depiction of DNA fragments used in shift assays. The corresponding primers are listed in Tab. S2.

**Fig. S7.**
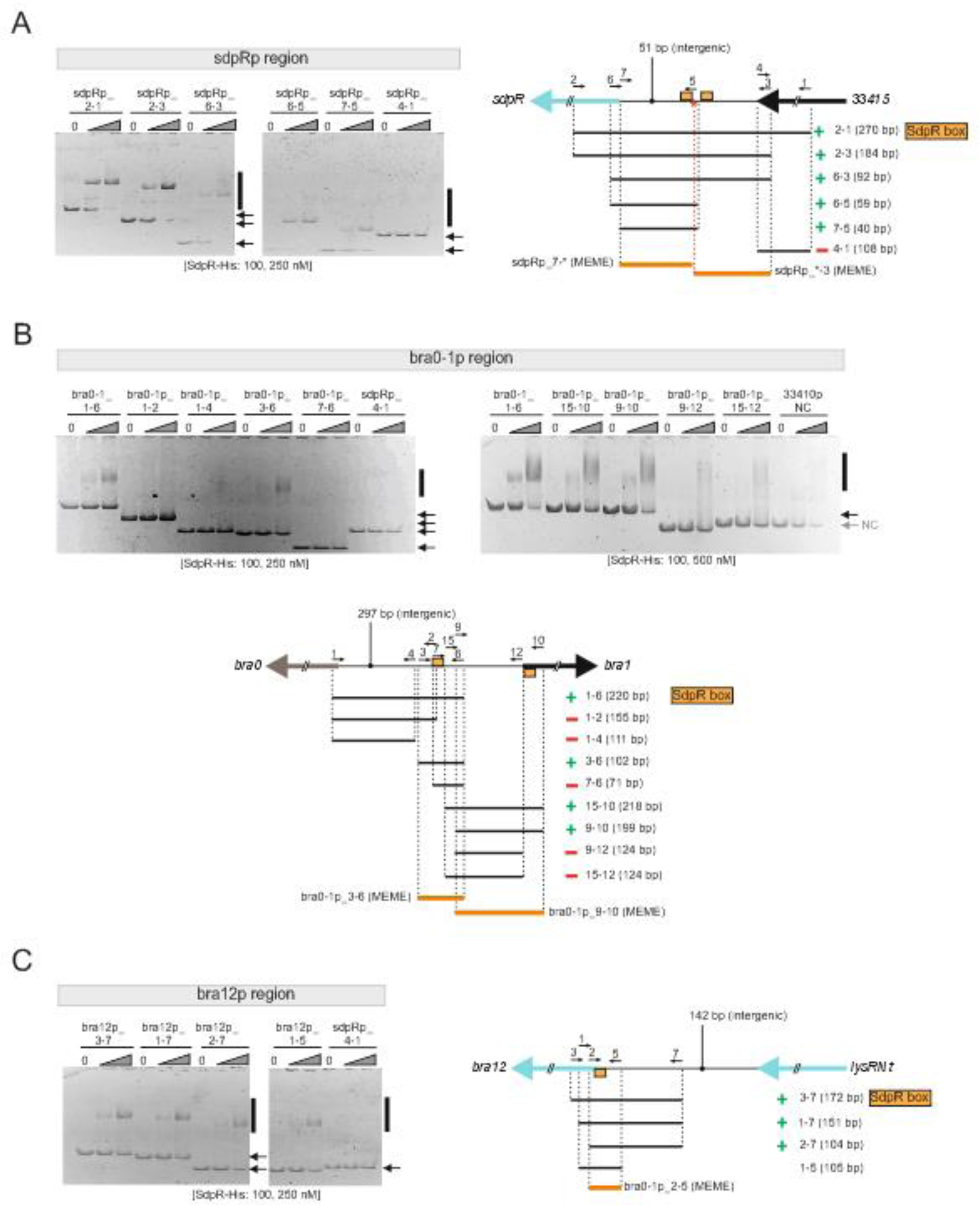
Detailed analysis of SdpR binding within promoter regions. Electrophoretic mobility shift assay (EMSA). The identification of SdpR binding sites within the *sdpR*, *bra0-1* and *bra12* promoter regions was shown in panels **(A)**, **(B)** and **(C)**, respectively. The recombinant SdpR-His protein was incubated with the subsets of PCR-amplified DNA fragments (primer list in Tab. S2). In the assays, constant amounts of unlabeled DNA and varying concentrations of the protein were used, as indicated. The *33140* gene promoter region (33410p) and sdpRp_4-1 fragments served as negative controls. Vertical black bars indicate protein-DNA complexes, and black and gray arrows indicate unbound DNA fragments. All panels contain a graphical depiction of the results. In these drawings, the green ‘**+**’ symbols indicate the interactions of SdpR-His with the corresponding DNA fragments, and the red symbols ‘**–**’ symbols represent the opposite observations, respectively. The regions used for *in silico* identification of binding sequences are marked with orange.

**Fig. S8.**
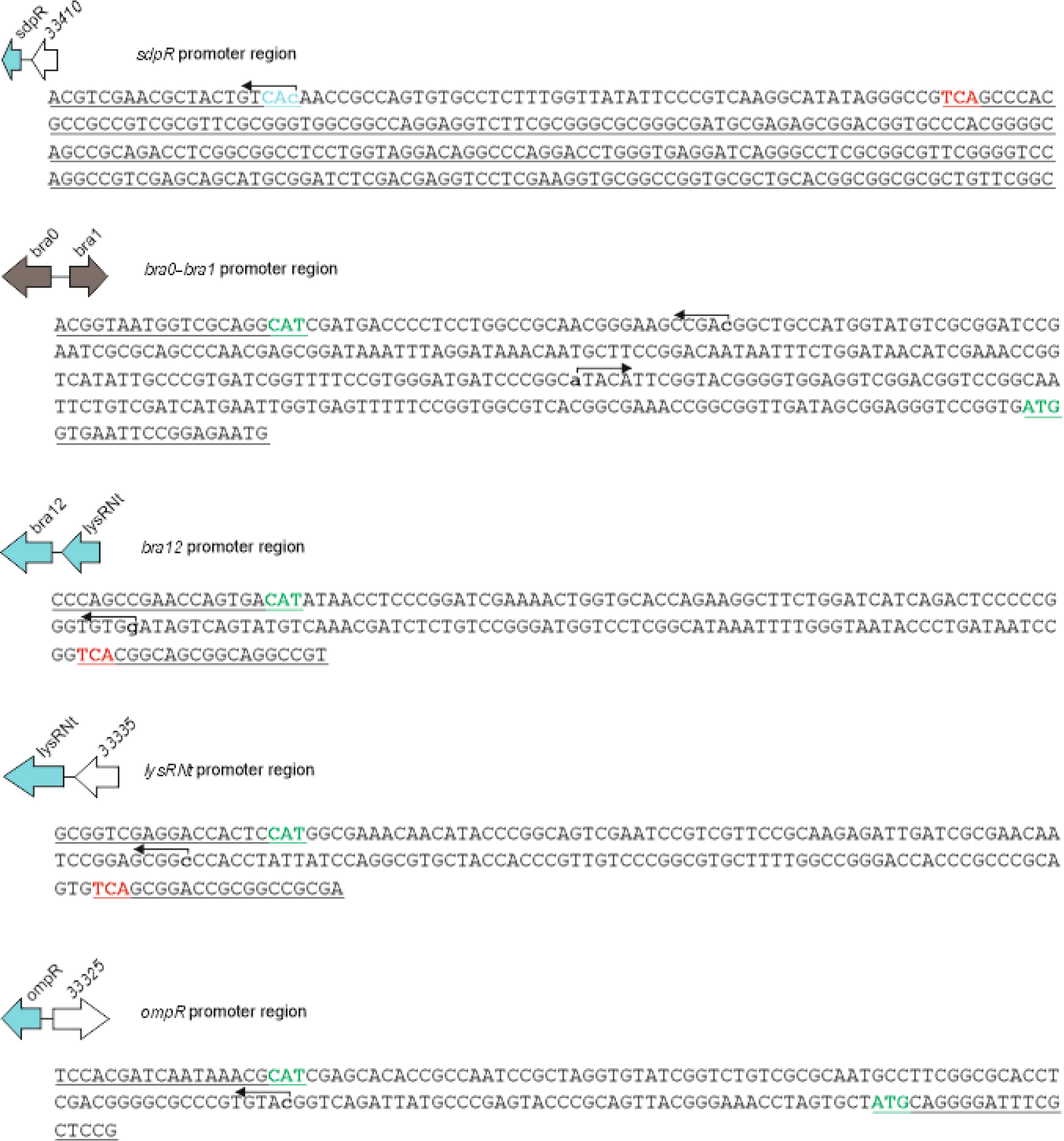
Transcription Start Sites (TSSs) in selected promoter regions within the Bra-BGC. RNA-seq. The TSSs are marked with bald lowercase letters and bent arrows. The first or the last twenty nucleotides of the gene sequences surrounding promoter regions are underlined; stop and start codons of corresponding genes are highlighted in red and green color, respectively. For original data, see Tab. S6.

**Fig. S9.**
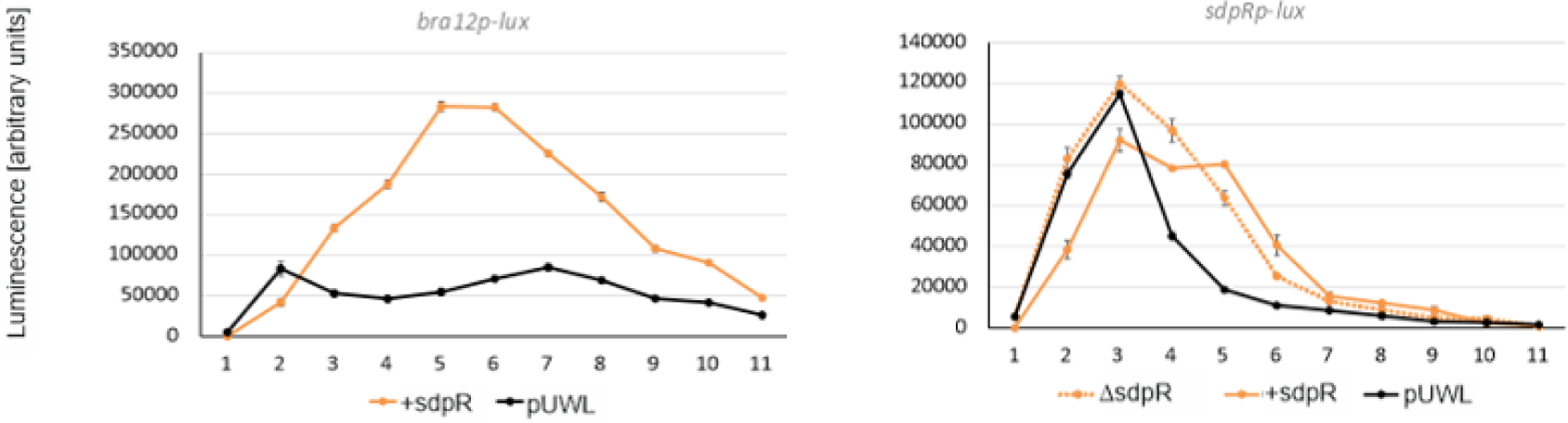
Impact of the SdpR regulator on promoter activities. Luciferase assays. To study the impact of SdpR overexpression, the measurements were conducted using *S. coelicolor* M1154 heterologous strains harboring *bra12p*, and *sdpRp* gene promoters delivered onto pFLUXH integrative plasmids, and replicating the pUWL201 plasmid overexpressing SdpR (+sdpR); the strains containing an empty pUWL201 plasmid served as the controls (pUWL). The impact of *sdpR* gene deletion was shown using a strain carrying the pFLUXH plasmid with *sdpRp,* and the bcaAB01_*Δ*sdpR fosmid (bottom panel). The luminescence readings were normalized against the OD_600_ of the corresponding cultures grown on solid DNA medium for 11 days.

**Fig. S10.**
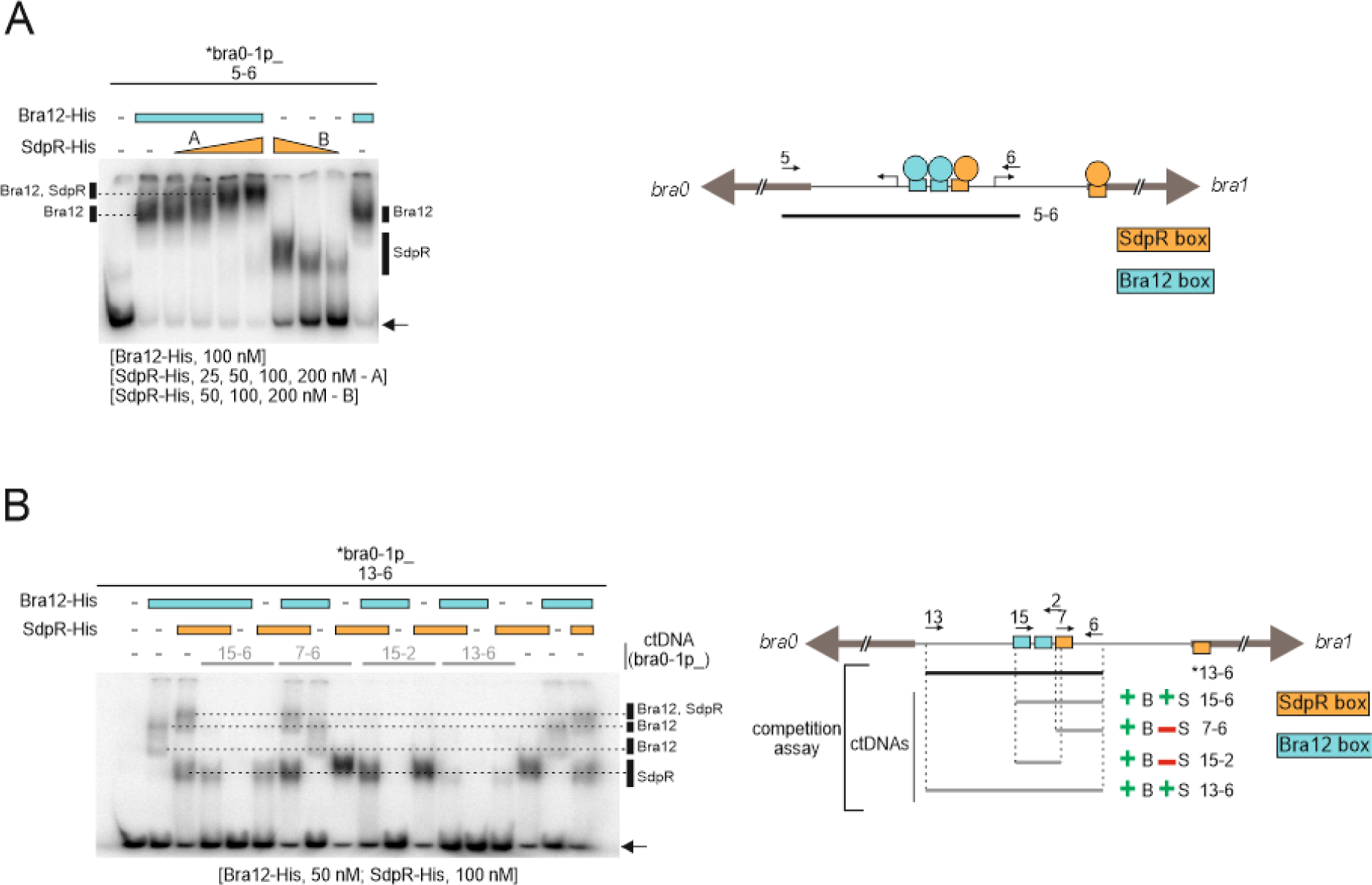
Simultaneous binding of Bra12 and SdpR to the bra0-1 intergenic region. **(A)** Electrophoretic mobility shift assays (EMSA). (left) The recombinant proteins (Bra12-His and SdpR-His) were incubated with the *bra0-1* intergenic region (^32^P-radiollabeled bra0-1p_5-6 fragment). A constant amount of DNA (∼ 5 mol) and the concentration of Bra12-His protein (blue bar) were used, and the SdpR-His protein was added at increasing concentrations (orange triangles), as indicated. Black vertical bars indicate protein-DNA complexes; a black arrow indicates unbound DNA. (right) Graphical summary of the EMSA and a model depicting simultaneous binding of Bra12 and SdpR to *bra0-1* intergenic region. The proteins are represented by coloured circles (blue: Bra12, orange: SdpR), the primers and the TSSs are shown with plain arrows and bent arrows, respectively. **(B)** competition EMSA. (left) The Bra12-His and SdpR-His at constant concentrations were incubated with the constant amount of the ^32^P-labeled DNA fragment (bra0-1p_13-6) comprising *bra0-1* intergenic region. The “cold-target DNA” competitors (ctDNA) were added to the reaction mixtures at 25 nM final concentration. Black vertical bars indicate protein-DNA complexes (marked also with dotted lines); a black arrow indicates unbound DNA. (right) Graphical summary of the assay. Radiolabeled DNA is indicated with asterisks (*). The green ‘**+**’ and the red ‘**–**’ symbols indicate the ability and lack of ability, respectively, of the corresponding ‘cold-target DNA’ fragment to outperform ^32^P-labeled DNA. Primers are indicated by short arrows. The numbers in brackets represent fragment sizes (bp).

**Fig. S11.**
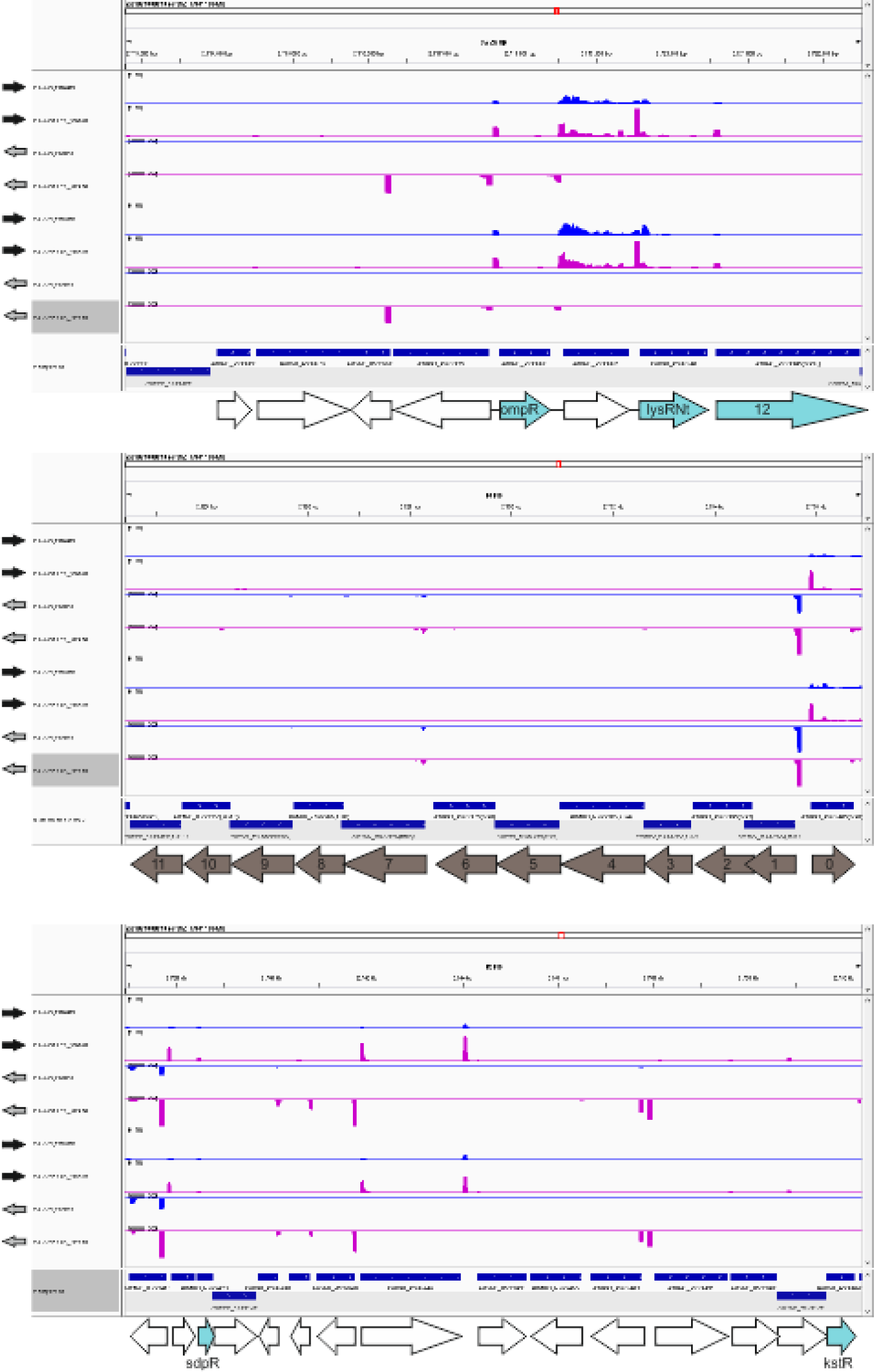
Expression tracks for the Bra-BGC. Raw gene expression data for *N. terpenica* IFM0406 grown for 33 and 48 hrs. Two replicates, R1 and R2, are shown. The expression tracks shown in blue and pink refer to normal RNA-seq and +TEX 5’-enriched libraries. The unit of the Y axis is coverage. The black and gray arrows on the left side show sense and anti-sense DNA strands of *N. terpenica* chromosome. Please note that due to software limitations, gene orientations are shown in a “natural” manner and appear on the chromosome sequence, and not the way we used throughout the article.

## Notes

### Competing Interest Statement

The authors have declared no competing interest.

